# Epigenetic Regulation of Nuclear Lamina-Associated Heterochromatin by HAT1 and the Acetylation of Newly Synthesized Histones

**DOI:** 10.1101/2021.06.28.450212

**Authors:** Liudmila V. Popova, Prabakaran Nagarajan, Callie M. Lovejoy, Benjamin D. Sunkel, Miranda L. Gardner, Meng Wang, Michael A. Freitas, Benjamin Z. Stanton, Mark R. Parthun

## Abstract

During S phase, eukaryotic cells must faithfully duplicate both the sequence of the genome and the regulatory information found in the epigenome. A central component of the epigenome is the pattern of histone post-translational modifications that play a critical role in the formation of specific chromatin states. During DNA replication, parental nucleosomes are disrupted and re-deposited on the nascent DNA near their original location to preserve the spatial memory of the epigenetic modifications. Newly synthesized histones must also be incorporated into the nascent chromatin to maintain nucleosome density. Transfer of modification patterns from parental histones to new histones is a fundamental step in epigenetic inheritance. Whether new histones play an active or passive role in epigenetic inheritance is unknown. Here we report that HAT1, which acetylates lysines 5 and 12 of newly synthesized histone H4 during replication-coupled chromatin assembly, regulates the epigenetic inheritance of chromatin states. HAT1 regulates the accessibility of large domains of heterochromatin termed HAT1-dependent Accessibility Domains (HADs). HADs are mega base-scale domains that comprise ~10% of the mouse genome. HAT1 functions as a global negative regulator of H3 K9me2/3 and HADs correspond to the regions of the genome that display HAT1-dependent increases in H3 K9me3 peak density. HADs display a high degree of overlap with a subset of Lamin-Associated Domains (LADs). HAT1 is required to maintain nuclear structure and integrity. These results indicate that HAT1 and the acetylation of newly synthesized histones are critical regulators of the epigenetic inheritance of heterochromatin and suggest a new mechanism for the epigenetic regulation of nuclear lamina-heterochromatin interactions.

## Introduction

When eukaryotic cells divide, not all of the information necessary for proper functioning of daughter cells is encoded in the primary DNA sequence of inherited genomes. Critical regulatory information is also relayed by patterns of chemical modifications to DNA bases and the histones packaging the genome, as well as in the 3-dimensional architecture of the genome. This regulatory information is inherited epigenetically during cell division. In mammals, the faithful transmission of epigenetic information is essential for life as it is critical for the control of cell proliferation, maintenance of cell identity and preservation of genome integrity (1–3).

The central process in cell division is genome duplication, which must occur with high fidelity and with accompanying inheritance of epigenetic information that is etched on DNA and chromatin. For example, patterns of CpG methylation are maintained following DNA replication by the maintenance DNA methyltransferase, DNMT1, which recognizes hemi-methylated DNA to restore CpG methylation patterns on daughter strands (4). By comparison, reproducing patterns of histone modifications on replicated chromatin or important aspects of 3-dimensional genomic structure are far more complex processes, and both are required for maintaining cell identity via proper regulation of transcriptional programs (5).

When a parental DNA duplex is replicated to produce two daughter duplexes, the chromatin packaging the DNA must be duplicated, as well. The replisome disrupts the parental nucleosomes in its path, leading to the dissociation of the histone octamer into an H3/H4 tetramer and two H2A/H2B dimers (6). The parental H3/H4 tetramer, which contains most of the epigenetically important modifications such as acetylation and methylation, is recycled onto one of the resulting daughter duplexes near its original location (7–14). This provides for the spatial memory of the parental histone modification patterns(15,16). These histone modification patterns play a central role in the formation of specific chromatin states by regulating inter-nucleosomal interactions and the association of non-histone proteins with chromatin.

As the parental H3/H4 tetramers are distributed to both daughter duplexes, maintaining correct nucleosome density requires that an equal number of newly synthesized H3/H4 tetramers be deposited on the newly replicated daughter duplexes (17). As a result, nascent chromatin is a 1:1 mixture of parental histones and newly synthesized histones. The accurate transfer of parental histone modification patterns to new histones is an essential step in the epigenetic inheritance of specific chromatin states(5,16).

The mechanisms underlying the transmission of parental histone modification patterns are best understood in the context of heterochromatin. Elegant work has established the “read-write” model to describe the transfer of transcriptionally repressive histone H3 methylation from parental to new histones(5,16,18,19). The lysine methyltransferases (KMTs) responsible for H3 K9me2/3 and H3 K27me3 also bind to, and are activated by, their cognate modification. For example, during the replication of constitutive heterochromatin, which is enriched in H3 K9me2 and H3 K9me3, the KMTs G9a and Suv39h1/2 can bind to the parental histones containing H3 K9me2/3. This brings the KMTs near the new histones and promotes the spread of the parental histone methylation pattern to neighboring new histones (5,20,21). While new histones are in close proximity to these KMTs in nascent chromatin, they acquire the parental patterns of methylation very slowly, suggesting that that this is a tightly regulated process(22,23).

Heterochromatic histone modification patterns also regulate genome architecture by regulating the interaction of chromatin with the structural components of the nucleus, such as the nuclear lamina(24–27). The nuclear lamina is a meshwork of intermediate filaments composed of A- and B-type lamins that coat the surface of the inner nuclear membrane through interactions with components of the nuclear membrane, such as the lamin B receptor (LBR) and Emerin(26,28). The regions of heterochromatin that interact with the nuclear lamina are known as LADs (lamin-associated domains). LADs range in size from ~0.1 Mb to 10 Mb, are gene poor and compose 30% to 40% of the mammalian genome(29,30). These domains are enriched in the repressive histone modifications H3 K9me2/3, H3 K27me3 and H4 K20me2/3. These modifications are essential for the interaction of LADs with the nuclear lamina, as loss of the lysine methyltransferases (KMTs) responsible for these methylation marks reduces heterochromatin-nuclear lamina association(31–33). Heterochromatin associated histone methylation promotes a direct physical association between heterochromatin and the nuclear lamina. Heterochromatin Protein 1 (HP1) is a reader for H3 K9me2/3. HP1 directly binds to both LBR and to PRR14, a proline-rich protein associated with the nuclear lamina through interactions with lamin A/C(24,34–36). In addition, LBR directly binds to H4 K20me2(37). Hence, direct physical interactions between heterochromatin and the nuclear lamina help drive the tethering of heterochromatin domains to the nuclear envelope and contribute to 3-dimensional genome architecture.

New histones have been considered as passive participants in epigenetic inheritance, serving as a blank canvas onto which the parental histone modification patterns are written (5,16). However, the newly synthesized H3 and H4 acquire specific patterns of modification prior to their deposition onto newly replicated DNA (17,38–41). The modification of new H3 begins with the co-translational monomethylation of K9 by ribosome-associated histone methyltransferase SetDB (42). H3 and H4 then form stable heterodimers in a chaperone-mediated process. The H3/H4 dimers then associate with the HAT1 complex that contains the histone acetyltransferase HAT1 and the histone chaperone Rbap46(43). HAT1 acetylates the new H4 on K5 and K12, an evolutionarily conserved modification pattern specific for newly synthesized molecules (38,44–47). Following nuclear import, the new H3 can also be acetylated, potentially by CBP or GCN5(48–59). The new H3/H4 complexes are then directed to sites of replication and deposited on newly replicated DNA by the chromatin assembly factor 1 (CAF1) complex (60–68). Following histone deposition, nascent chromatin assumes the correct 3- dimensional conformation during chromatin maturation. Chromatin maturation is a poorly characterized process that is accompanied by dynamic changes in the modification state of new histones. The predeposition patterns of acetylation are removed before the new histones acquire the post-translational modification patterns of neighboring parental histones (17). Whether the pre-deposition processing of newly synthesized H3 and H4 plays a regulatory role in the epigenetic inheritance of histone modification patterns is an open question.

HAT1 has emerged as a central regulator of new histone acetylation in mammalian cells. Studies in mouse embryonic fibroblasts (MEFs) have shown that, in addition to the expected acetylation of new H4 K5 and K12, the acetylation of new H3 is also HAT1-dependent. In particular, the deposition of H3 acetylated on K9 and 27 onto newly replicated DNA is lost in HAT1^−/−^ cells (38). Several lines of evidence suggest a link between HAT1 and regulation of the epigenetic inheritance of chromatin states. Complete knock out of HAT1 in mice results in a variety of developmental defects and neonatal lethality suggesting errors in cell fate decisions. In addition, HAT1^−/−^ MEFs display a high degree of genome instability(38). Finally, quantitative proteomics shows that HAT1 regulates the association of proteins with newly replicated DNA. Nascent chromatin assembled in the absence of HAT1 is enriched in factors involved in the formation of constitutive heterochromatin, including G9a, HP1, SMARCAD1, macro H2A, ATRX, KDM2A and the HELLS/CDCA7 complex(69).

We now report that HAT1 is a key regulator of the epigenetic inheritance of chromatin states and genome architecture. ATAC-Seq analysis of HAT1^+/+^ and HAT1^−/−^ MEFs demonstrates that HAT1 regulates the accessibility of specific chromosomal domains, which we have termed HAT1-dependent accessibility domains (HADs). HADs range in size from 0.1 to 10 Mb, are AT-rich, gene poor and heterochromatic. HAT1 functions as a global negative regulator of H3 K9me2 and K9me3, and HADs correspond to regions of the genome where HAT1 regulates the density of H3 K9me3. HADs display a high degree of overlap with Lamin-Associated Domains (LADs) suggesting that HAT1 and acetylation of newly synthesized histones regulate the association between heterochromatin and the nuclear lamina following DNA replication. We strengthen this link by reporting HAT1-dependent phenotypes consistent with disruption of nuclear lamina function. Finally, we propose a model for the epigenetic regulation of histone H3 methylation by HAT1 and newly synthesized histone acetylation and for the role of histone modification dynamics in the association of nascent chromatin with the nuclear lamina.

## Results

### Hat1 Regulates the Accessibility of Large Chromatin Domains

Nascent chromatin from HAT1^−/−^ cells is enriched for factors involved in constitutive heterochromatin formation, suggesting that the acetylation of newly synthesized histones regulates reestablishment of specific chromatin states following DNA replication. To test this hypothesis, we used ATAC-seq (Assay for Transposase Accessible Chromatin using sequencing) to profile genome-wide accessibility in Hat1^+/+^ and Hat1^−/−^ primary MEFs (70). The ATAC-Seq results demonstrate that HAT1 has a dramatic effect where 1859 sites were found to be differentially accessible (log_2_ fold change +/− 1, p<0.01), with 88% of these sites less accessible in HAT1^−/−^ cells (Figure 1A, Supplementary Table 1).

**Figure 1.**
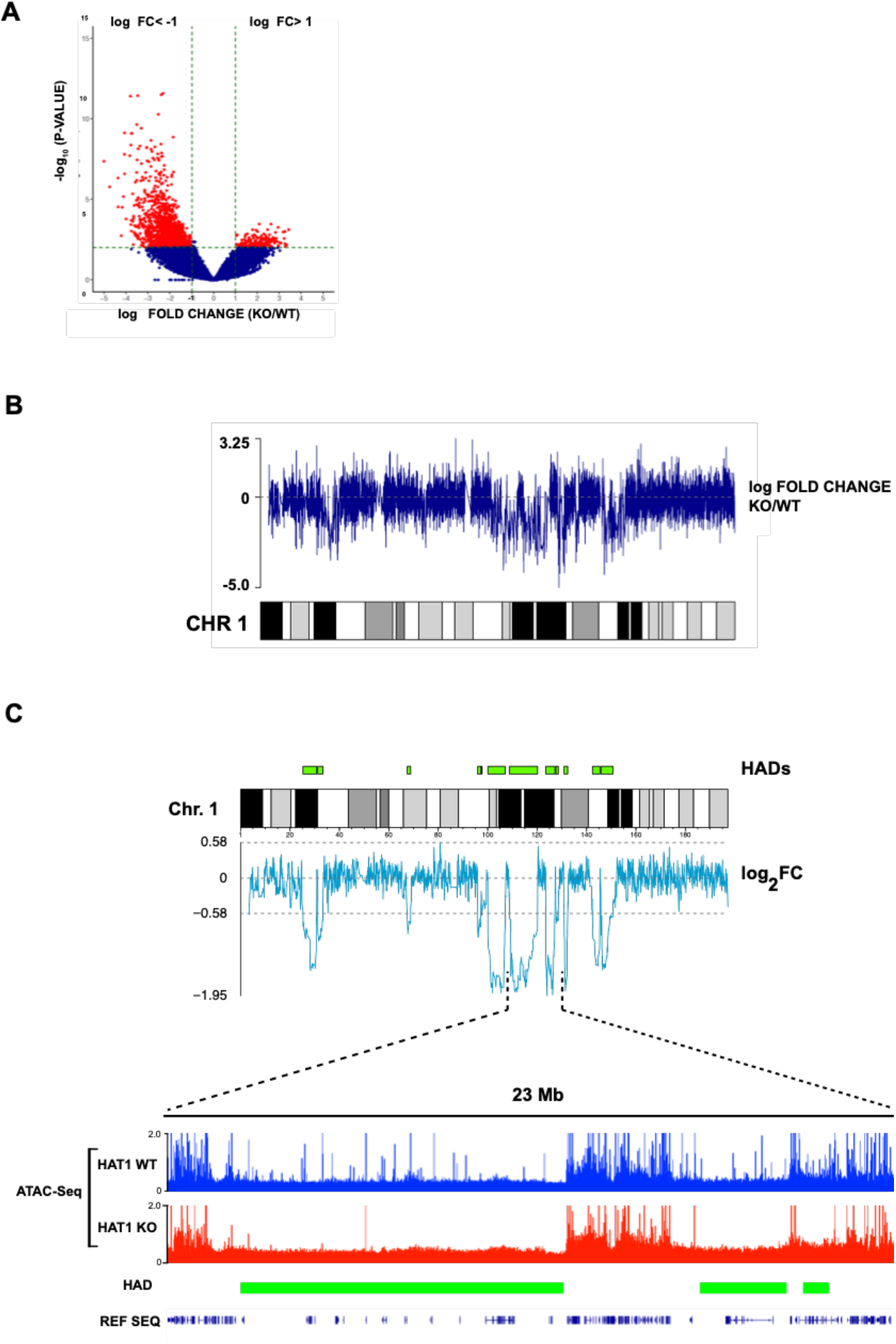
HAT1 regulates the accessibility of large chromatin domains. A, Volcano plot comparing ATAC-Seq peaks from HAT1^−/−^ vs. HAT1^+/+^ cells (log_2_ fold change +/− 1, p<0.01). B, The log_2_ fold change of ATAC-seq data from primary HAT1^+/+^ and HAT1^−/−^ MEFs plotted along mouse chromosome 1. C, (Top) The sliding window average (window size 25) of the log_2_ Fold Change (log_2_FC) of the ATAC-Seq signal from HAT1^−/−^ chromatin relative to HAT1^+/+^ chromatin is plotted below a karyoplot of chromosome 1. The location of HAT1-dependent accessibility domains (HADs) is indicated in green above the karyoplot. (Bottom) A genome browser view of the ATAC-Seq data from the indicated region of chromosome 1 (data is representative of biological triplicates). The location of HADs are indicated.

We localized the HAT1-dependent sites of accessibility relative to functional genomic features. The majority of HAT1-dependent sites of accessibility (57%) were in distal intergenic regions. The remaining sites were primarily in downstream introns (~31%) and promoter proximal regions (~8%) (Supplementary Figure 1A). HAT1-dependent sites of accessibility show a marked preference for low GC-content DNA, with more than 65% located in isochores L1 and L2 and 0 sites in isochore H3 (Supplementary Figure 1B). The HAT1-dependent sites of differential accessibility also display a preference for regions of low gene density (Supplementary Figure 1C).

Interestingly, the HAT1-dependent sites of accessibility are not evenly distributed throughout the genome. To visualize this distribution, we plotted the ATAC-Seq log_2_ fold change (log_2_ FC) difference between HAT1^+/+^ and HAT1^−/−^ cells across the genome. In these plots, values above 0 indicate increased accessibility in HAT1^−/−^ cells and values below 0 have decreased accessibility (Figure 1B). It is apparent that large domains of chromatin lose accessibility in HAT1^−/−^ cells. To computationally define these domains, we plotted a sliding window average of the log_2_ FC of the ATAC-Seq data (Figure 1C, Top). Regions where the log_2_ FC sliding window average is less than −0.58 (−1.5-fold change) were classified as <u>H</u>AT1-dependent <u>a</u>ccessibility <u>d</u>omains (HADs).

The bottom of Figure 1C shows a genome browser view of the ATAC-Seq data for a 23 Mb region of chromosome 1 that contains 3 HADs. The HADs are large domains that have characteristics of heterochromatin. HADs have lower gene density and lower accessibility than flanking euchromatic domains. Strikingly, the chromatin in HADs becomes almost entirely inaccessible in HAT1^−/−^ cells, while the flanking domains are unchanged. This suggests that the decrease in chromatin accessibility observed in the absence of HAT1 is largely due to decreased accessibility of heterochromatic regions rather than spreading of heterochromatin into neighboring euchromatic domains.

Across the genome (Y chromosome was excluded from analysis), we have identified 188 HADs that range in size from ~0.9 Kb to greater than 11 Mb, with ~48% of the HADs between 1 Mb and 5 Mb in size. In total, HADs encompass 10% of the mouse genome (Supplementary Table 2). The ATAC-Seq analysis was repeated with immortalized MEFs and highly similar patterns of HADs were observed, suggesting that the regulation of large domains of chromatin structure is a fundamental function of HAT1 (Supplementary Figure 2).

### HAT1 is a global Regulator of H3K9 Methylation

The decrease in constitutive heterochromatin accessibility upon HAT1 loss led us to hypothesize that HAT1 regulates the transmission of heterochromatic histone methylation patterns from parental histones to new histones. To test this hypothesis, we performed ChIP-Seq analysis of H3 K9me2 and H3 K9me3 in HAT1^+/+^ and HAT1^−/−^ MEFs. We used human chromatin as a spike-in control to permit a quantitative comparison of methylation levels between samples. Figure 2A shows a genome browser view of chromosome 1. There is a clear increase in the level of both H3 K9me2 and K9me3 in HAT1^−/−^ cells. Strikingly, the methylation increase is distributed across the length of the chromosome such that the overall patterns of H3 K9me2 and K9me3 are retained. Identical results are seen for all of the autosomes (data not shown). A global increase in H3 K9me2/3 is consistent with proteomic analyses that indicate a trend towards increased levels of H3 K9me2/3 and a decrease in the level of H3 K9me1 in HAT1^−/−^ MEFs (Supplementary Figure 3).

**Figure 2.**
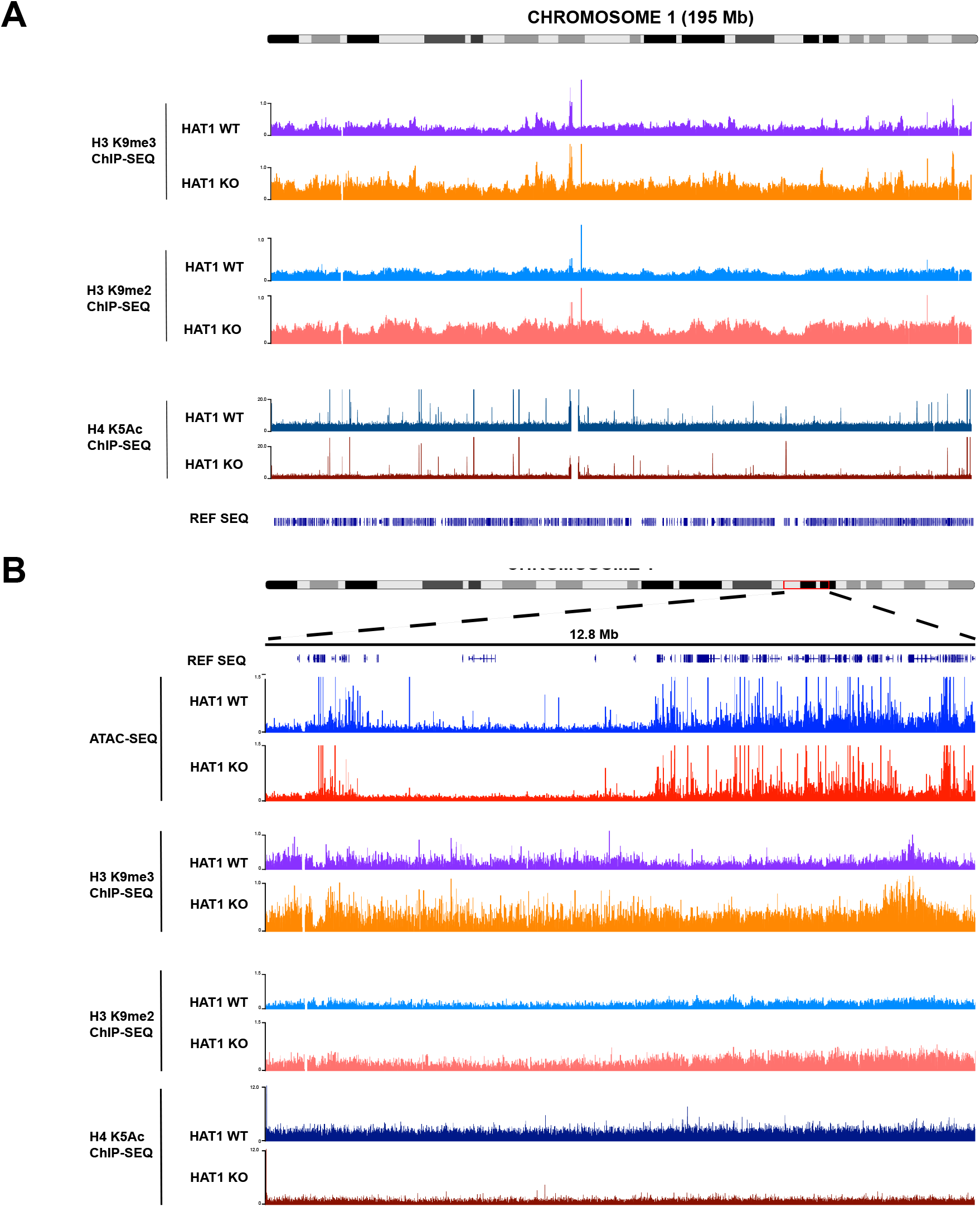
HAT1 is a global regulator of H3K9 methylation. A. Genome browser view of mouse chromosome 1 showing ChIP-Seq data of H3 K9me2, H3 K9me3 and H4 K5ac from immortalized HAT1^+/+^ and HAT1^−/−^ MEFs (as indicated). B. Genome browser view of a 12.8 Mb region of chromosome 1 showing ATAC-Seq data and ChIP-Seq data of H3 K9me2, H3 K9me3 and H4 K5ac from immortalized HAT1^+/+^ and HAT1^−/−^ MEFs (as indicated).

The H3 K9me2/3 ChIP-Seq data suggests that HAT1-dependent regulation of H3 K9 methylation occurs uniformly across the genome. A genome-wide role for HAT1 is consistent with previous reports demonstrating that nearly 100% of newly synthesized histone H4 is diacetylated on K5 and K12(39,47,71). To obtain additional support for a genome-wide function for HAT1, we analyzed the effect of HAT1 loss on H4 K5 acetylation. This analysis is complicated by the transient nature of the acetylation of new H4, as this modification is lost during chromatin maturation(72,73). We predicted that ChIP-Seq analysis of H4 K5 acetylation from an asynchronous culture would reveal a basal level of acetylation that is due to the random distribution of active replication fork progression across the genome. Regions of HAT1-dependent new H4 acetylation should be indicated by a decrease in this basal level of K5 acetylation. As seen in Figure 6A, ChIP-Seq analysis of H4 K5 acetylation in HAT1^+/+^ and HAT1^−/−^ cells indicates that there is a uniform decrease of basal H4 K5 acetylation across the entirety of chromosome 1. Hence, these results suggest that HAT1 is responsible for the acetylation of new histones that are deposited throughout the genome and that HAT1 and the acetylation of new histones function as global negative regulators of H3 K9 methylation.

The decrease in accessibility in HADs was not strictly correlated to increased levels of H3 K9me2/me3. Figure 2B shows a genome browser view of a 12.8 Mb region of chromosome 1. The levels of H3 K9 me2/3 are elevated in the regions of HAT1-dependent accessibility. However, there were also increased levels of H3 K9 methylation in flanking regions whose accessibility does not change. In addition, there is a uniform decrease of H4 K5Ac across regions where accessibility is both HAT1-dependent and HAT1-independent. Hence, HAT1-dependent alterations in chromatin accessibility do not appear to be linked to the overall levels of H3 K9 methylation.

### HAT1 regulates H3 K9me3 density in HADs

When analyzed by population-based methods such as ChIP-Seq, the concentration of H3 K9 methylation in a given region of chromatin is indicated by two measurements; the intensity of the ChIP-Seq signal and the density of the ChIP-Seq signal. As HADs do not correlate with HAT1-dependent changes in H3 K9me2/3 intensity, we analyzed HAT1-dependent changes in H3 K9 methylation peak density. In Figure 3A, the plot above the karyoplot of chromosome 1 depicts the change in the density of H3 K9me32peaks between HAT1^+/+^ and HAT1^−/−^ cells, where values above zero reflect an increase in peak density in the HAT1^−/−^ cells. Loss of HAT1 has little effect on the density of H3 K9me2 peaks. However, HAT1 has a significant effect on H3 K9me3 peak density across chromosome 1 (Figure 3B). We observed a striking correlation between HADs and the regions of the chromosome that display increases in H3 K9me3 peak density. Similar results are observed across all autosomes (data not shown). These data indicate that chromatin accessibility in HADs is linked to the HAT1-dependent regulation of H3 K9me3 density.

**Figure 3.**
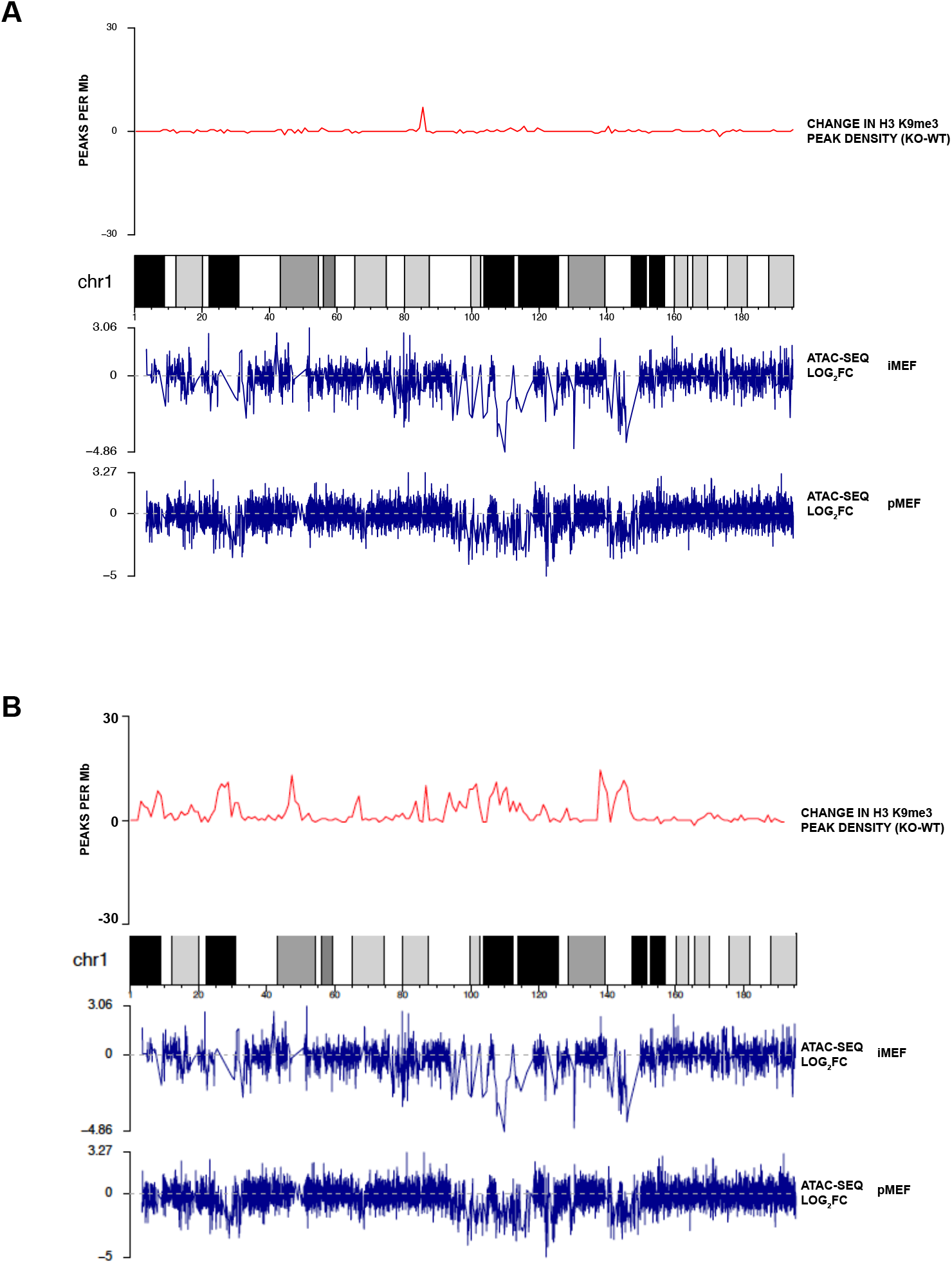
HAT1 regulates H3 K9me3 peak density in HADs. **A.** The change in the density of H3 K9me2 peaks (peaks/Mb), determined by comparing ChIP-Seq data from HAT1^+/+^ and HAT1^−/−^ MEFs, is plotted above a karyoplot of mouse chromosome 1. Below the karyoplot are plots of the log_2_ FC of ATAC-Seq data between HAT1^+/+^ and HAT1^−/−^ immortalized and primary MEFs (as indicated). B. The change in the density of H3 K9me3 peaks (peaks/Mb), determined by comparing ChIP-Seq data from HAT1^+/+^ and HAT1^−/−^ MEFs, is plotted above a karyoplot of mouse chromosome 1. Below the karyoplot are plots of the log_2_ FC of ATAC-Seq data between HAT1^+/+^ and HAT1^−/−^ immortalized and primary MEFs (as indicated).

### HAT1-dependent chromatin accessibility is not tightly linked to transcription

It was recently shown that in regions of euchromatin, the ground state of nascent chromatin following DNA replication is inaccessibility. Accessibility is acquired during chromatin maturation as genes become transcriptionally active(74). To determine whether the loss of accessibility in HAT1^−/−^ cells is the result of a loss of transcription, we performed RNA-Seq analysis in HAT1^+/+^ and HAT1^−/−^ MEFs. Consistent with the overall decrease in accessibility of HAT1^−/−^ chromatin, the primary effect of HAT1 loss was a decrease in transcription (Figure 4A). The expression of 709 genes decreased by at least 1.5-fold and 283 genes increased by at least 1.5-fold (p-value<0.05). However, comparison of the ATAC-Seq and RNA-Seq data suggests that the HAT1-dependent changes in chromatin accessibility are not tightly linked to changes in transcription. Of the 709 genes down-regulated in primary HAT1^−/−^ MEFs, only 39 (~5%) are located in HADs. Consistent with their heterochromatic nature and low gene density, most HADs have low levels of transcription. For example, Figure 4B shows a HAD from chromosome 16. Within the HAD there is very little transcription and there is no difference between the HAT1^+/+^ and HAT1^−/−^ cells. This raises the interesting possibility that in the wake of DNA replication, the reemergence of the correct patterns of accessibility in heterochromatic HADs occurs through mechanisms distinct from those governing euchromatic inheritance.

**Figure 4.**
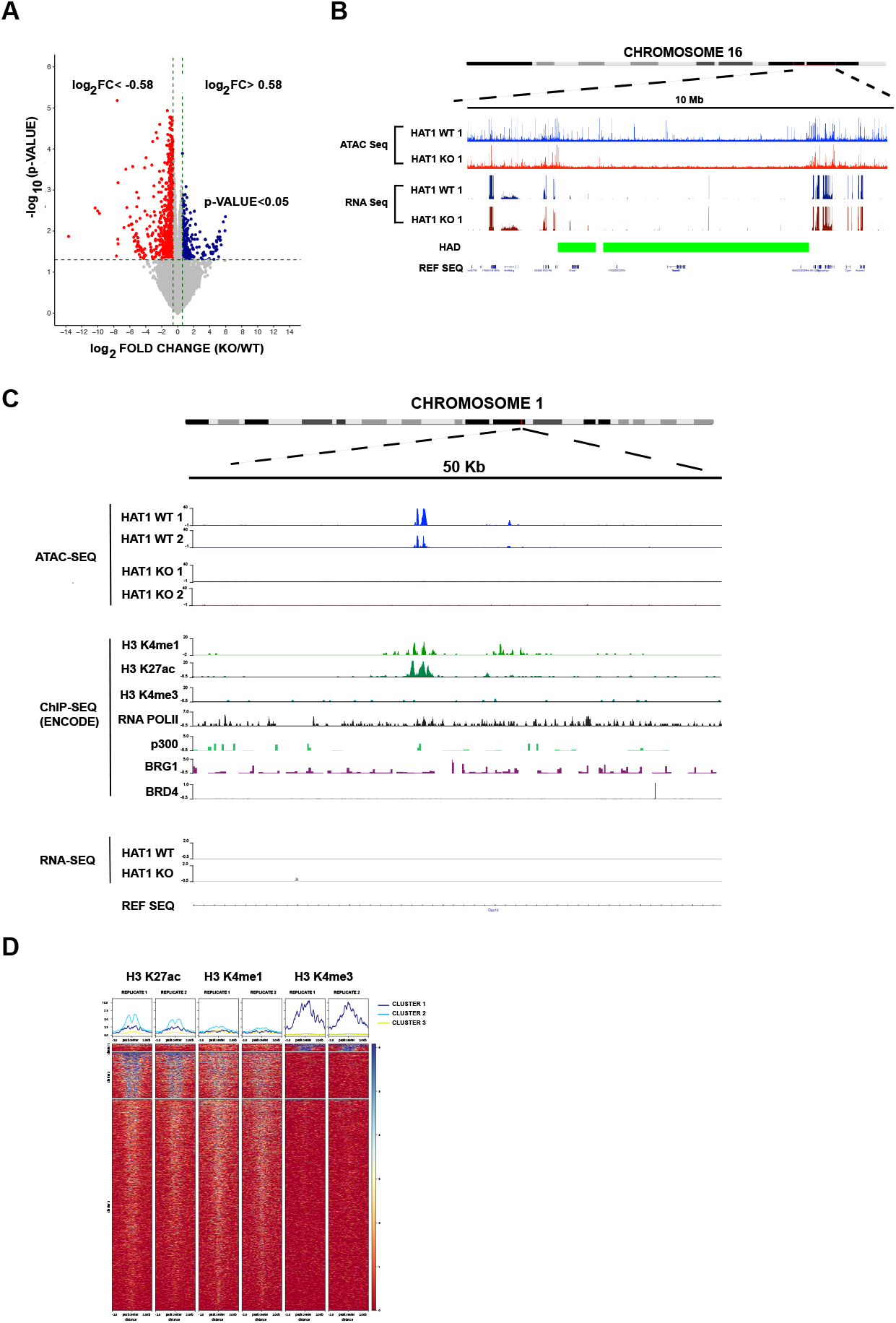
HAT1-dependent chromatin accessibility is not tightly linked to transcription. A. Volcano plot of RNA-Seq data from primary HAT1^+/+^ and HAT1^−/−^ MEFs. B. Genome browser view of a 10 Mb region of chromosome 16 showing ATAC-Seq and RNA-Seq data from primary HAT1^+/+^ and HAT1^−/−^ MEFs. Location of HADs is indicated by the green boxes. C. Genome browser view of a 50 Kb region of chromosome 1 showing the indicated tracks. ATAC-Seq and RNA-Seq was generated by our laboratory in HAT1^+/+^ and HAT1^−/−^ MEFs. ChIP-Seq data was obtained from MEFs by the ENCODE project. D. The 1859 HAT1-dependent sites of accessibility were aligned with publicly available ChIP-Seq data for H3 K27ac, H3 K4me1 and H3 K4me3. The sites were divided into 3 groups following unsupervised hierarchical clustering. Heatmaps show the abundance of each modification relative to the centers of the peaks of accessibility +/− 2Kb. E. Genome browser view of a 22.7 Mb segment of mouse chromosome 1

### HAT1-dependent sites of accessibility have characteristics of active enhancers

To better understand the nature of the HAT1-dependent sites of accessibility embedded within heterochromatin, we compared our ATAC-Seq data to publicly available ChIP-Seq datasets for histone post-translational modifications and chromatin modifying proteins. The HAT1-dependent sites of accessibility have properties of active enhancers as there is a significant correlation with H3 K27ac and H3 K4me1 (Figure 4C and 4D). However, only a small subset of these sites are also associated with H3 K4me3 peaks, consistent with the lack of connection between HAT1-dependent accessibility and active transcription.

### HADs localize to lamin-associated domains

The nuclear lamina is an important structural component of the nucleus that provides a framework for the 3-dimensional architecture of the genome. The association of LADs with the nuclear lamina localizes these domains to the periphery and the heterochromatin compartment of the nucleus(25,26,75,76). LADs range in size from 0.1 Mb to 10 Mb, tend to be gene poor and are enriched in the L1 and L2 isochores. The physical characteristics of HADs are similar to those of LADs (25,26). We compared our ATAC-Seq data with available lamin B1 localization in MEFs determined by DamID and found a remarkable degree of overlap (Figure 5A). In the karyoplots shown in Figure 5B, it is clear that HADs are predominantly located within LADs throughout the genome (86% of HADs overlap with LADs). These data indicate that HAT1 regulates the accessibility of large chromosomal domains associated with the nuclear lamina.

**Figure 5.**
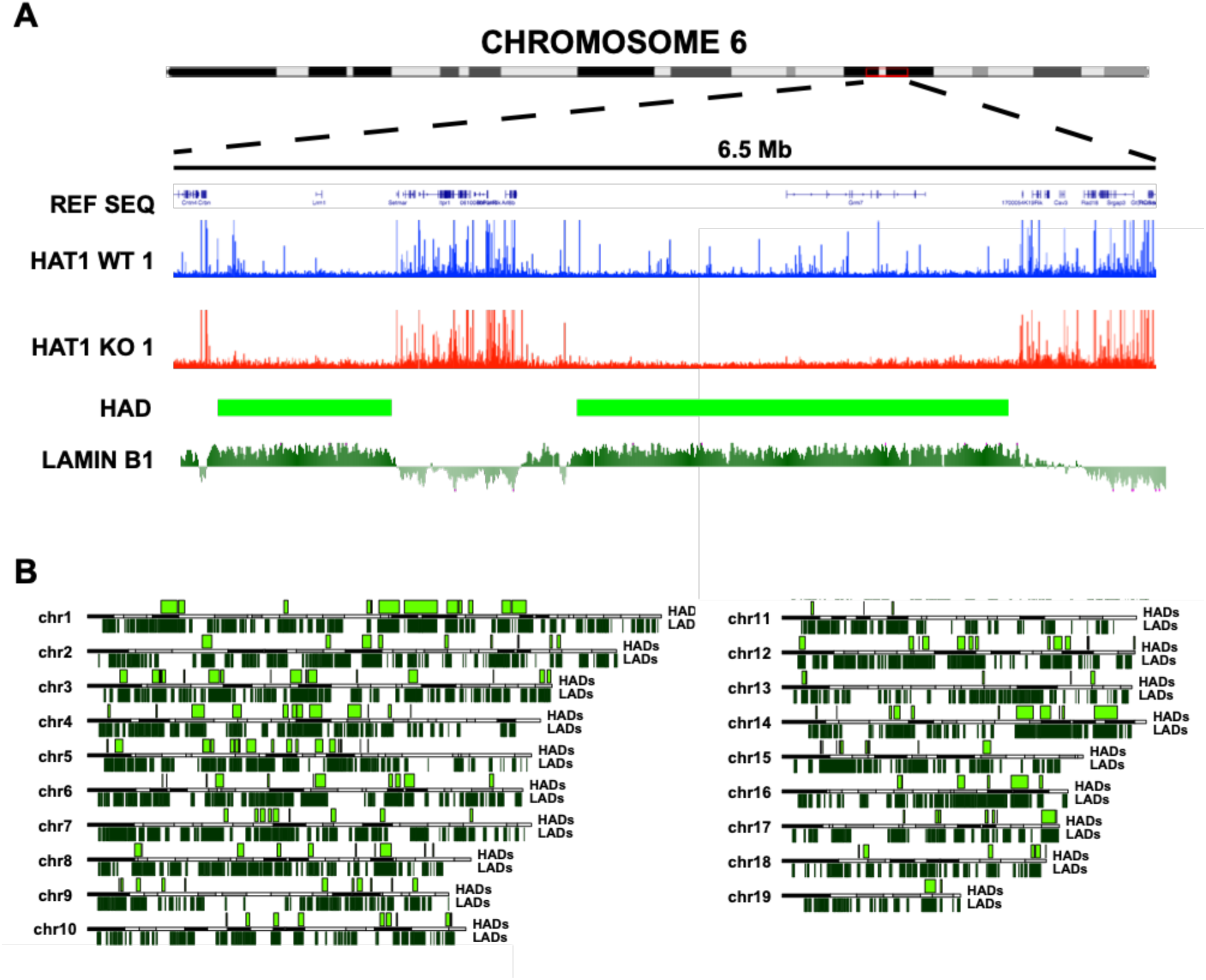
HADs display a high degree of overlap with LADs. A. Genome browser visualization of ATAC Seq data from HAT1^+/+^ (blue) and HAT1^−/−^ (red) cell lines. For the indicated region of chromosome 6. Lamin B1 DamID data (dark green) from MEFs was obtained from the UCSC browser(97). B. Karyoplots of each autosome with location of HADs on the top and LADs on the bottom.

### HAT1 is required to maintain nuclear structure and stability

Previous studies have suggested a link between HAT1 and nuclear lamina function. For example, complete knockout of HAT1 in mice results in neonatal lethality with a number of developmental phenotypes that are very similar to those seen following the complete knockout of the LMNB1 gene, including hyper-proliferation of lung mesenchymal cells and craniofacial bone defects(38,77). In addition, senescence of MEFs is highly sensitive to the level of HAT1(78). Finally, mice heterozygous for HAT1 have a dramatically shortened lifespan and develop multiple signs of early onset aging(78).

The nuclear lamina is a critical determinant of nuclear morphology. To determine whether HAT1 is functionally linked to the nuclear lamina, we examined several aspects of nuclear structure and stability. We measured nuclear size in HAT1^+/+^ and HAT1^−/−^ MEFs and found that loss of HAT1 leads to a significant increase in nuclear area (Figure 6A). This is consistent with a number of studies that have demonstrated an increase in nuclear size associated with decreases in the expression of lamins or with mutations that compromise lamina function(79,80).

**Figure 6.**
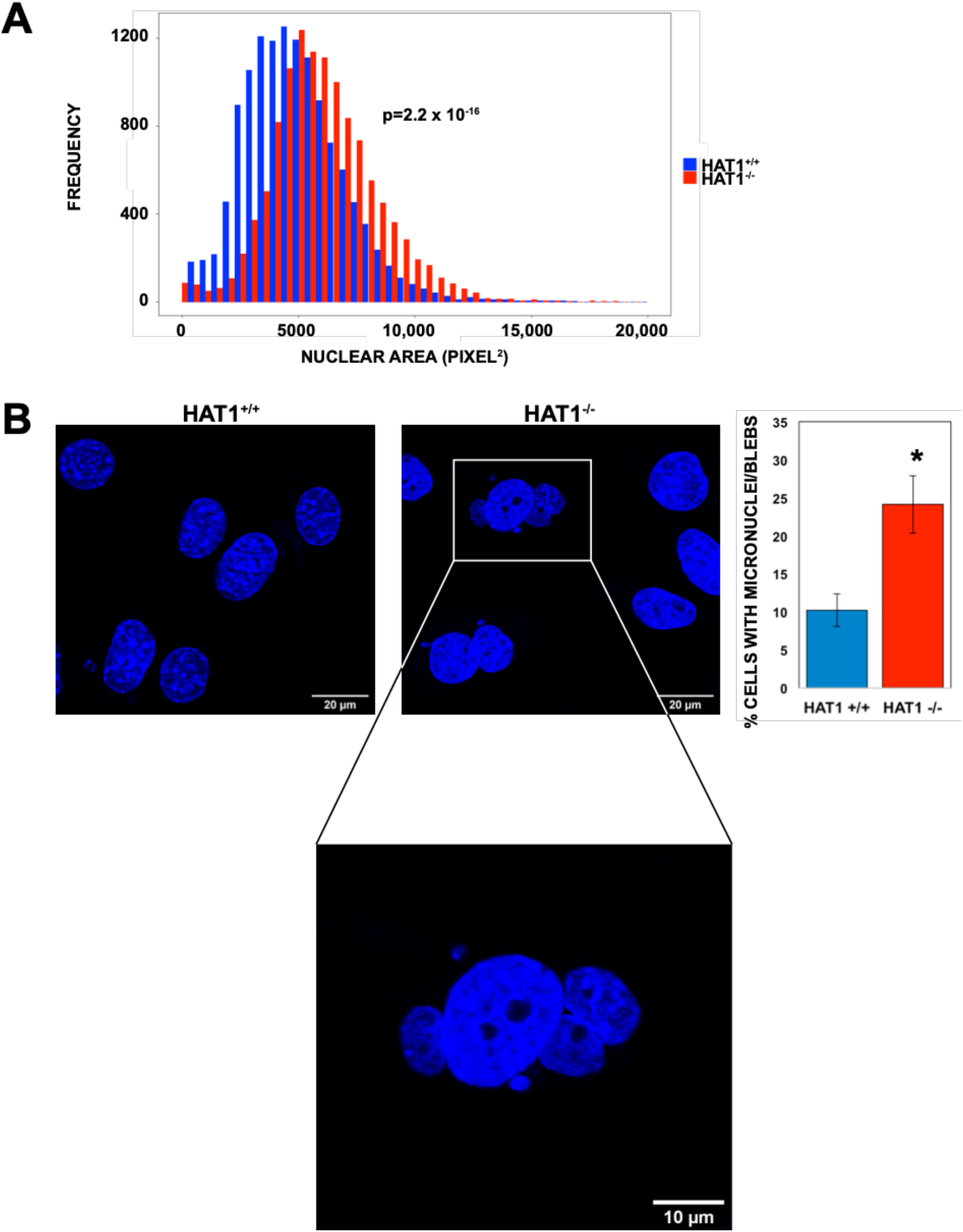
HAT1 is required to maintain nuclear structure and stability. A. HAT1^+/+^ (n=12,980) and HAT1^−/−^ (n=11,796) cells were stained with DAPI, imaged and nuclear area determined with Image J. B. Cells of the indicated genotype were visualized with DAPI. The percentage of cells containing nuclear blebs or micronuclei (marked by white arrows) was quantitated (n= 1207 and 894 cells for HAT1^+/+^ and HAT1^−/−^, respectively, *= p-value <0.05).

Defects in the nuclear lamina can also compromise the structural integrity of the nucleus. An interesting example is related to the protein acetyltransferase Mof. Analysis of Mof-dependent acetylation identified multiple sites of acetylation in the lamins, particularly lamin A. This acetylation is functionally relevant as mutations in Mof or the Mof-dependent sites of acetylation in lamin A cause increased nuclear blebbing and formation of micronuclei (81). We identified a similar phenotype in nuclei from HAT1^−/−^ cells, as there is a significant increase in the frequency of nuclear blebbing and micronuclei relative to that observed in control HAT1^+/+^ cells (Figure 6B). We conclude that HAT1 deficient cells display an array of phenotypes similar to those seen in cells with defects in the nuclear lamina further supporting that HAT1 is functionally linked to nuclear lamia -associated heterochromatin.

Our results demonstrate that HAT1 plays a central role in the epigenetic inheritance of chromatin states. Specifically, we demonstrate that HAT1 regulates the accessibility of large domains of chromatin that we have termed HADs (<u>H</u>AT1-dependent <u>A</u>ccessibility <u>D</u>omains). In addition, HAT1 functions as a global negative regulator of H3 K9 methylation and chromatin accessibility in HADs is regulated by HAT1-dependent regulation of H3 K9me3 density. HADs are megabase-scale domains that coincide with heterochromatic regions of chromatin that interact with the nuclear lamina (LADs). HAT1 is directly linked to the nuclear lamina as HAT1 loss leads to defects in nuclear structure and integrity.

## Discussion

In the wake of a replication fork, nascent chromatin contains a 1:1 mixture of parental and newly synthesized histones. The presence of H3 K9me2/3 on parental histones provides the template for the propagation of these modifications to new histones and the epigenetic inheritance of constitutive heterochromatin structure (15). The ‘read-write” model for this epigenetic inheritance of heterochromatin posits that parental H3 K9me2/me3 serve as binding sites for the KMTs G9a and Suv39h1/2. This recruits these KMTs to nascent chromatin and into the proximity of neighboring newly synthesized histones for the propagation of the methylation marks(5,16,82). Here, we test the hypothesis that HAT1 and the acetylation of the newly synthesized histones play an active regulatory roles in the epigenetic inheritance of histone H3 K9 methylation and constitutive heterochromatin.

We propose that HAT1 and the acetylation of new histones serve as a spatial and/or temporal brake on the transfer of H3 K9me2/3 from parental histones to new histones (Figure 7A). This transient inhibition of the propagation of K3 K9me2/3 would provide an opportunity for the ordered and regulated reestablishment of constitutive heterochromatin structure following DNA replication. A regulatory role for new histone acetylation is consistent with a number of previous reports. Early studies involving the treatment of cells in S phase with histone deacetylase inhibitors indicated that a delay in the removal of new histone acetylation prevented the generation of stable and mature chromatin(72). More recent studies have used proteomics to characterize the acquisition of H3 K9me2/3 by newly synthesized histones. Surprisingly, methylation of H3K9 occurs with slow kinetics following deposition of newly synthesized histones on nascent chromatin, with new histones taking an entire cell cycle to acquire the parental level of methylation(22,23).

**Figure 7.**
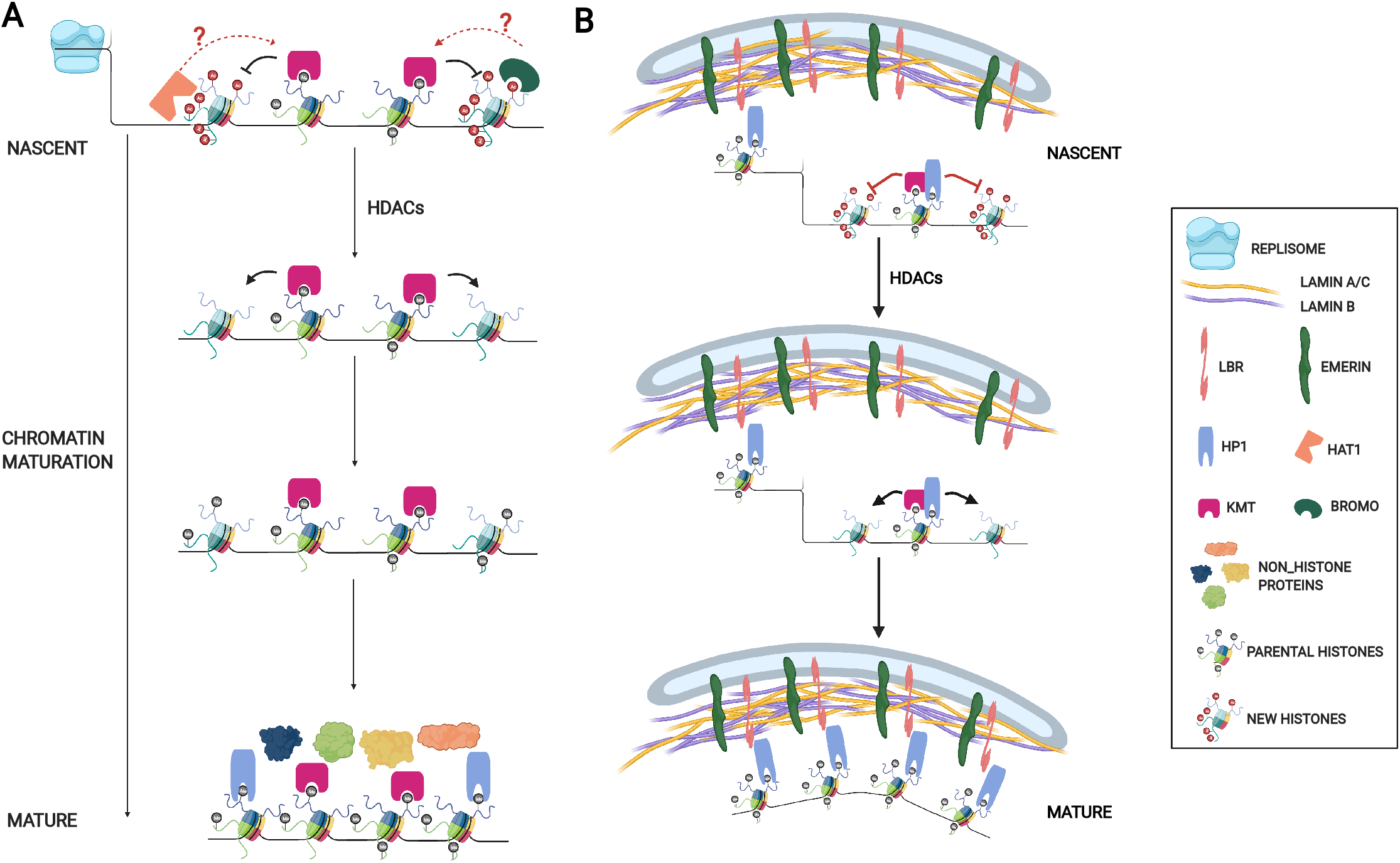
Model of epigenetic regulation of nuclear lamina-associated heterochromatin domains by HAT1 and the acetylation of newly synthesized histones. A. Schematic diagram showing the potential role of HAT1 and the acetylation of newly synthesized histones in the epigenetic inheritance of histone H3 methylation in heterochromatin. B. Schematic diagram describing the regulation of nuclear lamina-heterochromatin interactions following DNA replication by nascent chromatin maturation.

There are multiple, non-mutually exclusive, mechanisms by which HAT1 and the acetylation of newly synthesized histones may regulate the epigenetic inheritance of histone methylation patterns on nascent chromatin (Figure 7). HAT1 is required for the acetylation of newly synthesized H3 and H4 and for the deposition of H3 and H4 with specific sites of acetylation onto newly replicating DNA. In particular, H3 acetylated on K9 and K27 and H4 acetylated on K5 and K12 are absent on nascent chromatin from HAT1^−/−^ cells(38). The deposition of new H3/H4 tetramers containing H3 K9ac would directly block the ability of KMTs to methylate these new histones (Figure 7A). An important caveat of this mechanism is that a relatively small proportion of new H3 appears to be acetylated on K9. Pulse labeling with ^14^C-lysine demonstrated that ~3% of new H3 molecules were acetylated on K9(71). Hence, the direct inhibition of H3 K9me2/3 by K9 acetylation is only likely to play a significant regulatory role if new H3 containing this modification is specifically targeted and concentrated at discreet regions of the genome, such as HADs. In this context, it is interesting to note that increasing H3 K27ac can disrupt the interaction of LADs with the nuclear lamina(83,84).

Contrary to new H3, essentially all new molecules of H4 are diacetylated on K5 and K12 by HAT1(39,71). Our results indicate that HAT1-modified new H4 is uniformly distributed throughout the genome. Hence, the genome-wide effect on H3 K9me2/3 is more likely linked to HAT1 or the acetylation of new H4. The acetylation of new H4 creates binding sites for bromodomain proteins on nascent chromatin and it was recently shown that association of Brg1, BRD3 and Baz1a with nascent chromatin is HAT1-dependent(69). The presence of these proteins on nascent chromatin may directly block the spread of H3 K9 methylation or may function to promote the localized assembly of open chromatin (Figure 7A). The latter possibility is consistent with the active enhancer-like structure of the HAT1-dependent sites of accessibility in constitutive heterochromatin, given the central role of bromodomain proteins in the formation of active enhancers in the context of euchromatin(85).

The transient physical association of HAT1 with newly replicated DNA opens up the possibility that HAT1 is directly involved in the regulation of H3 K9me2/3 dynamics on nascent chromatin (Figure 7A)(38,69,86). The presence of HAT1 on nascent chromatin may sterically block the activity of KMTs, as the RBBP7/HAT2 subunit of the HAT1 complex also binds the NH_2_-terminal tail of histone H3. Alternatively, HAT1 may negatively regulate H3 K9 methylation directly or indirectly through its acetyltransferase activity. Indeed, a recent report identified the HAT1-dependent acetylation of a number of chromatin modifying proteins, including KMT5b(87). HAT1 was also recently proposed to play a central role in the regulation of nuclear acetyl-CoA availability and the absence of HAT1 may decrease heterochromatin accessibility through decreased acetyl-CoA availability(88).

In addition to genome-wide increases in H3 K9me2/3, loss of HAT1 also results in the formation of megabase scale domains of highly inaccessible chromatin (HADs). A link between these two phenomena is suggested by the correlation between HADs and HAT1-dependent changes in the density of H3 K9me3 peaks. Recent reports have suggested that constitutive heterochromatin structure is formed by liquid-liquid phase separation or polymer-polymer phase separation(89–93). An important aspect of these models is that phase separation is mediated by the concentration and valence of HP1 interactions with chromatin. In the absence of HAT1, regions of the genome where the density of H3 K9me3 increases would provide an environment with a high local HP1 concentration and the opportunity for multi-valent interactions. This may push the HP1 concentration past the normal threshold and drive the assembly of an aberrantly condensed form of heterochromatin.

In the context of euchromatin, accessibility following DNA replication develops as a consequence of transcriptional restart (74). Chromatin accessibility in regions of constitutive heterochromatin may develop through a distinct mechanism. Comparison of our ATAC-Seq and RNA-Seq data indicates that heterochromatic chromatin accessibility is not strictly linked to proximal transcription. However, the sites of accessibility possess properties similar to those of active enhancers in euchromatin. This raises the question of the function of sites of chromatin accessibility in gene poor regions of heterochromatin. These sites of accessibility may be important for facilitating long-range chromatin interactions that are important for proper 3-dimensional genome architecture. These sites of accessibility may also be involved in creating regional chromatin environments that allow for the interaction of heterochromatin components with structural components of the nucleus, such as the nuclear lamina.

The nuclear lamina is a critical structure that provides the framework for the 3- dimensional organization of the genome. Our understanding of nuclear lamina formation and its association with specific chromatin domains is based largely on studies of its assembly following mitosis(25–27). However, it is likely that passage of a replication fork through lamin-associated chromatin disrupts lamin-chromatin interactions. The timely reestablishment of the association between nascent chromatin and the nuclear lamina would then be essential for the epigenetic inheritance of global genome architecture.

We propose a model for the epigenetic regulation of nuclear lamina-heterochromatin interactions following DNA replication (Figure 7B). The replication fork disrupts the nuclear lamina-heterochromatin interaction as the parental nucleosomes are displaced. The recycled parental histones carry H3 K9me2/3, which is necessary for reestablishing contact with the nuclear lamina through factors such as LBR and PRR14, which directly interact with HP1(24,29,33,35,36). We propose that the association of newly synthesized histones with the nuclear lamina is regulated by dynamic changes in their modification state. The interaction between the new histones and the nuclear lamina would develop during chromatin maturation as the new histones acquire H3 K9me2/3. Hence, DNA replication and chromatin maturation provide a window of opportunity to epigenetically regulate global genome architecture through modulating the localization of heterochromatin with the nuclear periphery.

## Acknowledgements

We would like to thank Dr. aaron Goldman, Dr. Moray Campbell, Dr. Paul Wade and Dr. Gene Oltz for helpful discussions. This work was supported by funding from the National Institutes of Health (GM062970 to MRP and CMBP T32 to LVP), the St. Baldrick’s Foundation (Berry Strong Fund award to BZS), the Mark Foundation for Cancer Research (ASPIRE Award to BZS), Andrew McDonough B+ Foundation (Childhood Cancer Research Grant to BZS), and the CancerFree KIDS Foundation (New Idea Award to BDS). Microscopy facilities were supported by grants P30 NS104177 and S10 OD02684. Proteomics facilities were supported by P30 016058 to the Ohio State University Comprehensive Cancer Center.

## Author Contributions

L.P. designed and performed epigenomic experiments, performed bioinformatic analysis of epigenomic and proteomic studies, and helped write the manuscript. P.N. generated cell lines, performed RNA-Seq, H4 K5ac ChIP-Seq and epiprofiling, analyzed data, and edited the manuscript. C.M.L. performed analysis of nuclear structure and integrity assays, analyzed data and edited the manuscript. B.D.S. performed epigenomic experiments. M.L.G. performed and analyzed the mass spectrometric analysis of histones and edited the manuscript. M.W. performed bioinformatic analyses of epigenomic data. M.A.F. designed experiments and analyzed experimental data. B.D.S. designed experiments, analyzed data and edited the manuscript. M.R.P. designed experiments, analyzed data, and wrote the manuscript.

## Materials and Methods

### MEF Cell Culture

e12.5 to e14.5 embryos were dissected from the pregnant female and internal organs were removed. The embryonic tissue was disaggregated using an 18-gauge syringe and brought to single cell suspension with Trypsin incubation at 37°C. Cells were then plated onto 100mm tissue culture plates, passaged upon confluency and maintained in Dulbecco’s modified Eagle medium (DMEM-Sigma) with 10% fetal bovine serum (FBS-Gibco) and 1X Pen/Strep antibiotics (Sigma). Genomic DNA was isolated from fibroblasts by standard methods using phenol: chloroform isolation and genotypes were confirmed by PCR using the following pairs of primers P1: 5′-GCC TGG TGA GAT GGC TTA AAC -3′ and P2: 5′-GCA AGT AGT ATG ACA AGA GGT AGG =3′. PCR was performed under following conditions; 95°C for 50 min followed by 29 cycles at 95°C for 40 sec., 54.6°C for 30 sec. and 72°C for 60 sec. and final extension for 5 min. at 72°C. The WT and mutant alleles yielded product sizes of 916 bp and 478 bp respectively. SV40 T immortalized MEFs (iMEFs) were derived from primary HAT1^+/+^ and HAT1^−/−^ embryonic day 13.5 embryos. Early passage primary MEFs were transformed with SV-40 T antigen containing plasmid pBSSVD2005 (ADDGENE, Cambridge, MA) to generate immortalized MEFs. Early passage cells were seeded at 25% confluency in six-well plates and transfected with 2 ug of expression vector using Fugene reagent (Roche). Cells were harvested and seeded into 100 mm dishes after 48 h of transfection. The cells were split at 1 in 10 dilutions until passage 5. For the ATAC-seq and ChIP-seq experiments, cells were grown in DMEM media (Sigma) supplemented with 10% FBS (Sigma) and 1X Penicillin/Streptomycin antibiotics (Gibco).

### Data Analysis and Code Availability

All custom data analysis, unless otherwise indicated, was performed using R version 3.6.3 (2020-02-29). The chromosome plots were generated using karyoploteR package version 1.12.4 in R (94). Additional plots were generated using *ggplot2* (95).All custom code generated for this publication is available as supplementary material.

### pMEF ATAC-Seq Library Preparation

Primary MEFs isolated from HAT1^+/+^ (n= 3) and HAT1^−/−^ (n=3) embryos (5*10^4 cells) were collected, resuspended in cold PBS, and centrifuged for 5min at 500g at 4°C. Next, cells were washed with 1mL cold RSB buffer (10mM Tris-HCl, pH 7.4, 10 mM NaCl, and 3mM MgCl_2_). Cells were centrifuged at 500g for 10 min at 4°C, resuspended in 250μL of freshly made Lysis Buffer (250μL of RSB buffer and 2.5μL of 10% NP-40), and incubated for 5 minutes on ice. Cells were centrifuged at 500g for 10 min at 4°C. Isolated nuclei were resuspended in 50μL of the transposition mix (22.5μL of H2O, 25μL of the 2X Tagment DNA Buffer, 2.5μL of Tagment DNA Enzyme (both Illumina)) and incubated at 37°C for 30 min. Transposition reactions were purified with the MinElute PCR Purification Kit (Qiagen) and eluted in EB buffer. ATAC-seq libraries were amplified with Nextera primers (Illumina) by PCR and purified with AMPure beads (Beckman Coulter). Resulting ATAC libraries were sequenced with paired-end reads.

### pMEF ATAC-Seq Data Analysis

ATAC-seq reads were processed with the ENCODE ATAC-sequencing pipeline version 1.6.1. The reads were aligned to the mouse genome version mm10 using bowtie2, and the peaks were called using MACS2. Differential accessibility analysis was carried out in R using the DiffBind package version 2.14.0.

### Genomic Features Annotation and Gene Density Analysis

The annotation of the ATAC-seq differentially occupied peaks was carried out using the ChIPseeker package (version 1.22.1) in R. Gene density analysis was performed using the TxDb.Mmusculus.UCSC.mm10.knownGene package (version 3.10.0) in R.

### Isochore Analysis

Dataset, containing mouse isochore genomic coordinates and isochore classification based on the G/C content was taken from Costantini et al.(96). The genomic coordinates of the isochores were converted from mm9 to mm10 genome assemblies using the UCSC liftOver tool. Percentages of differentially occupied sites in different isochores were calculated using the plyranges package (version 1.6.10) in R.

### Computational Identification of HADs

To identify HADs computationally, sliding window averages of the ATAC-seq HAT1^−/−^ to HAT1^+/+^ log2 fold change were calculated using slider (version 0.1.5) package in R. Next, a custom R algorithm (available as supplementary material) was used to systematically identify HADs. With this algorithm, any two adjacent regions that fell below the set cutoff of −0.58 log_2_ fold change or 1.5 fold change and had no more than 1Mb of gap between themselves were considered to be a part of the same HAD. As a result, 189 HADs were identified. 1 HAD localized on chromosome Y was removed from further analysis.

### HADs and LADs Data Analysis

MEF LADs dataset was downloaded from Peric-Hupkes et al.(97). The genomic coordinates of the LADs were converted from mm9 to mm10 genome assemblies using the UCSC liftOver tool. Overlap between HADs and LADs was calculated using the plyranges package (version 1.6.10) in R.

### RNA-Seq

RNA-seq was performed as previously described in Nagarajan et al. (98).

### ENCODE ChIP-seq Data Analysis

Bigwig files for H3K4me1, H3K4me3, and H3K27ac histone marks in MEFs were downloaded from ENCODE (datasets ENCSR000CAZ, ENCSR000CBA, and ENCSR000CDI respectively). Heatmaps were generated using deeptools version 3.5.1.

### imMEF ATAC-Seq Library Preparation and Data Analysis

Immortalized MEFs isolated from Hat1^+/+^ (n= 2) and Hat1^−/−^ (n=2) (1*10^5 cells) were collected; nuclei were isolated from the cells and subjected to a transposition reaction with Tn5 transposase by Novogene. Sequencing libraries were amplified by PCR and sequenced with paired-end reads. Raw reads were aligned to the mouse genome version mm10 using bwa version 0.7.12 and peaks were called using MACS2 version 2.1.0. Differential occupancy analysis was performed using DESeq2 in R.

### Immunoblot analysis of HAT1 mutant MEFs

For immunoblot anlaysis, protein lysates from MEFs were prepared by using RIPA buffer (100 mM Tris-HCl pH 7.4; 300 mM NaCl; 2% NP-40; 1% sodium Deoxycholate; 0.2% SDS). Protein lysates were separated in 12% acrylamide gels and transferred to nitrocellulose blotting membrane (cat.# 10600004-GE Healthcare Life Sciences). The membrane was blocked with 5% skim milk in TBS-T (20 mM Tris-HCl at pH 7.4, 150 mM NaCl, 0.1% Tween-20) for an hour at room temperature and then incubated with primary antibodies against HAT1 (Abcam, ab12163), histone H3 (Abcam, ab1791), histone H4 (Abcam, ab10158), histone H3 K9me2 (Abcam, ab1220), H3 K9me3 (Abcam, ab8898) and alpha Tubulin (abcam ab7291) incubated overnight at 4°C. HRP-conjugated secondary antibodies and Pierce ECL or Brighstar Femto chemiluminescent ( Pierce cat.# 32106 ; Brightstar cat. # XR94) were used for detection.

### Nuclear Structure and Integrity Assays

Three HAT1+/+ or HAT1−/− MEF cell lines were seeded in equal quantities on coverslips and allowed to attach for 24 h. Cells were then permeabilized with 0.5% Triton X-100 and fixed with 4% PFA simultaneously for 15 min, rinsed with PBS, and fixed again with 4% PFA for 10 min at room temperature. After several PBS washes cells were blocked with 5% BSA for 1 h at room temperature. BSA was removed with PBS. Nuclei were stained with 20 mM Hoechst 33342 Fluorescent Stain and mounted on slides using Vectashield. Images were acquired using a Zeiss LSM 900 Airyscan 2 Point Scanning Confocal microscope and Zen Blue 3.0. Quantification was completed using ImageJ version 1.52t by identifying nuclear regions, measuring the size, and recording nuclei that exhibited nuclear abnormalities.

### H3K9me2 and H3K9me3 ChIP

Immortalized MEFs isolated from HAT1^+/+^ (n= 2) and HAT1^−/−^ (n=2) embryos were collected, washed with PBS, and resuspended in CiA fixing buffer (50 mM HEPES, pH 8, 1 mM EDTA, 100 mM NaCI, and 0.5 mM EGTA). Cells were fixed with 1% formaldehyde (Thermo Scientific) for 10 min at room temperature. Crosslinking reaction was quenched with 0.125M glycine. Cells were washed with PBS supplemented with 1X of Complete protease inhibitor cocktail (Roche), pelleted and frozen. The pellets were thawed and resuspended 0.8mL of ice-cold TE buffer (pH 8.0) supplemented with 1X of Complete protease inhibitor cocktail (Roche). Human Dbt immortalized myoblasts were added to the mouse cells to be used as a spike-in control. Samples were sonicated with an Active Motif sonicator for 20 min at 30% amplitude using a 30sec on/off cycle. Sheared chromatin samples were adjusted to RIPA 200 mM NaCl buffer (final concentrations of 0.1% SDS, 0.1% sodium deoxycholate, 1% Triton X-100, and 200 mM NaCl in 1X TE). Samples were incubated on ice for 5 minutes and centrifuged at 13,000 rpm for 10 min at 4°C. H3K9me2 and H3K9me3 antibodies (Abcam ab1220 and ab8898, respectively) were added to the supernatant, and the samples were incubated at 4°C with rotation for 2 hours. Next, pre-washed with RIPA 200mM NaCl buffer Dynabeads (Invitrogen) were added to the samples for overnight incubation. After incubation, Dynabeads were isolated and washed twice with RIPA buffer (0.1% SDS, 0.1% sodium deoxycholate, and 1% Triton X-100 in 1X TE), twice with RIPA 200 mM NaCl Buffer, twice with LiCl Buffer (250 mM LiCl, 0.5% NP-40, 0.5% sodium deoxycholate in 1X TE), and twice with TE buffer. Beads were isolated, sonicated DNA was decrosslinked overnight and eluted using the MinElute PCR kit (Qiagen).

### H3K9me2 and H3K9me3 ChIP-Seq Library Preparation and Data Analysis

Sequencing libraries were prepared by PCR amplification and sequenced with paired-end reads by Novogene. Reads were aligned to the mouse genome version mm10 using BWA version 0.7.12, and the peaks were called using MACS2 version 2.1.0. Differential accessibility analysis was carried out using the DiffBind package. ChIP-seq signal density was calculated using the karyoploteR package version 1.12.4 in R(94).

### H4 K5ac ChIP-Seq

Mouse embryonic fibroblasts were fixed with 1% formaldehyde in flasks at room temperature for 10 min and subsequently washed with ice-cold PBS before a 0.1-M glycine solution was added to stop the fixation. After chromatin fragmented to 200 to 500 bp by sonication using Bioruptor. chromatin templates from 20 million cells were used for ChIP experiment. Samples were immunoprecipitated with 2-4 μg of H4K5 antibody overnight at 4 °C followed by washing steps. After reverse crosslinking, the ChIP DNA fragment were purified. After reverse crosslinking, the ChIP DNA fragment were purified and repaired followed by treatment with Taq polymerase to generate single-base 3’ overhangs used for adaptor ligation. Libraries were prepared with the Illumina ChIP-seq Sample Prep Kit and sequenced using a genome analyzer (Illumina). ChIP-seq reads were mapped to the most recent mouse genome (mm9) using IGV (Integrative Genomics Viewer) and only the uniquely mapping reads were used for further analysis.

## Notes

### Competing Interest Statement

The authors have declared no competing interest.

### Summary of Updates

The revised version contains additional data and analysis.

## References

1. Ishiuchi, T., Enriquez-Gasca, R., Mizutani, E., Bošković, A., Ziegler-Birling, C., Rodriguez-Terrones, D., Wakayama, T., Vaquerizas, J.M. and Torres-Padilla, M.E. (2015) Early embryonic-like cells are induced by downregulating replication-dependent chromatin assembly. Nat Struct Mol Biol, 22, 662–671.

2. Yadav, T., Quivy, J.P. and Almouzni, G. (2018) Chromatin plasticity: A versatile landscape that underlies cell fate and identity. Science, 361, 1332–1336.

3. Cheloufi, S., Elling, U., Hopfgartner, B., Jung, Y.L., Murn, J., Ninova, M., Hubmann, M., Badeaux, A.I., Euong Ang, C., Tenen, D. et al. (2015) The histone chaperone CAF-1 safeguards somatic cell identity. Nature, 528, 218–224.

4. Bronner, C., Alhosin, M., Hamiche, A. and Mousli, M. (2019) Coordinated Dialogue between UHRF1 and DNMT1 to Ensure Faithful Inheritance of Methylated DNA Patterns. Genes (Basel), 10.

5. Stewart-Morgan, K.R., Petryk, N. and Groth, A. (2020) Chromatin replication and epigenetic cell memory. Nat Cell Biol, 22, 361–371.

6. Annunziato, A.T. (2005) Split decision: what happens to nucleosomes during DNA replication? J Biol Chem, 280, 12065–12068.

7. Liu, S., Xu, Z., Leng, H., Zheng, P., Yang, J., Chen, K., Feng, J. and Li, Q. (2017) RPA binds histone H3-H4 and functions in DNA replication-coupled nucleosome assembly. Science, 355, 415–420.

8. Bellelli, R., Belan, O., Pye, V.E., Clement, C., Maslen, S.L., Skehel, J.M., Cherepanov, P., Almouzni, G. and Boulton, S.J. (2018) POLE3-POLE4 Is a Histone H3-H4 Chaperone that Maintains Chromatin Integrity during DNA Replication. Mol Cell, 72, 112–126.e115.

9. Clément, C. and Almouzni, G. (2015) MCM2 binding to histones H3-H4 and ASF1 supports a tetramer-to-dimer model for histone inheritance at the replication fork. Nat Struct Mol Biol, 22, 587–589.

10. Huang, H., Strømme, C.B., Saredi, G., Hödl, M., Strandsby, A., González-Aguilera, C., Chen, S., Groth, A. and Patel, D.J. (2015) A unique binding mode enables MCM2 to chaperone histones H3-H4 at replication forks. Nat Struct Mol Biol, 22, 618–626.

11. Richet, N., Liu, D., Legrand, P., Velours, C., Corpet, A., Gaubert, A., Bakail, M., Moal-Raisin, G., Guerois, R., Compper, C. et al. (2015) Structural insight into how the human helicase subunit MCM2 may act as a histone chaperone together with ASF1 at the replication fork. Nucleic Acids Res, 43, 1905–1917.

12. Wang, H., Wang, M., Yang, N. and Xu, R.M. (2015) Structure of the quaternary complex of histone H3-H4 heterodimer with chaperone ASF1 and the replicative helicase subunit MCM2. Protein Cell, 6, 693–697.

13. Groth, A., Corpet, A., Cook, A.J., Roche, D., Bartek, J., Lukas, J. and Almouzni, G. (2007) Regulation of replication fork progression through histone supply and demand. Science, 318, 1928–1931.

14. Foltman, M., Evrin, C., De Piccoli, G., Jones, R.C., Edmondson, R.D., Katou, Y., Nakato, R., Shirahige, K. and Labib, K. (2013) Eukaryotic replisome components cooperate to process histones during chromosome replication. Cell Rep, 3, 892–904.

15. Reveron-Gomez, N., Gonzalez-Aguilera, C., Stewart-Morgan, K.R., Petryk, N., Flury, V., Graziano, S., Johansen, J.V., Jakobsen, J.S., Alabert, C. and Groth, A. (2018) Accurate Recycling of Parental Histones Reproduces the Histone Modification Landscape during DNA Replication. Mol Cell, 72, 239–249 e235.

16. Escobar, T.M., Loyola, A. and Reinberg, D. (2021) Parental nucleosome segregation and the inheritance of cellular identity. Nat Rev Genet, 22, 379–392.

17. Annunziato, A.T. (2012) Assembling chromatin: The long and winding road. Biochim Biophys Acta, 1819, 196–210.

18. Alabert, C., Loos, C., Voelker-Albert, M., Graziano, S., Forné, I., Reveron-Gomez, N., Schuh, L., Hasenauer, J., Marr, C., Imhof, A. et al. (2020) Domain Model Explains Propagation Dynamics and Stability of Histone H3K27 and H3K36 Methylation Landscapes. Cell Rep, 30, 1223–1234.e1228.

19. Allshire, R.C. and Madhani, H.D. (2018) Ten principles of heterochromatin formation and function. Nat Rev Mol Cell Biol, 19, 229–244.

20. Hathaway, Nathaniel A., Bell, O., Hodges, C., Miller, Erik L., Neel, Dana S. and Crabtree, Gerald R. (2012) Dynamics and Memory of Heterochromatin in Living Cells. Cell, 149, 1447–1460.

21. Reinberg, D. and Vales, L.D. (2018) Chromatin domains rich in inheritance. Science, 361, 33–34.

22. Scharf, A.N., Barth, T.K. and Imhof, A. (2009) Establishment of histone modifications after chromatin assembly. Nucleic Acids Res, 37, 5032–5040.

23. Alabert, C., Barth, T.K., Reveron-Gomez, N., Sidoli, S., Schmidt, A., Jensen, O.N., Imhof, A. and Groth, A. (2015) Two distinct modes for propagation of histone PTMs across the cell cycle. Genes Dev, 29, 585–590.

24. Harr, J.C., Gonzalez‐Sandoval, A. and Gasser, S.M. (2016) Histones and histone modifications in perinuclear chromatin anchoring: from yeast to man. EMBO reports, 17, 139–155.

25. van Steensel, B. and Belmont, A.S. (2017) Lamina-Associated Domains: Links with Chromosome Architecture, Heterochromatin, and Gene Repression. Cell, 169, 780–791.

26. Briand, N. and Collas, P. (2020) Lamina-associated domains: peripheral matters and internal affairs. Genome Biol, 21, 85.

27. Hoskins, V.E., Smith, K. and Reddy, K.L. (2021) The shifting shape of genomes: dynamics of heterochromatin interactions at the nuclear lamina. Curr Opin Genet Dev, 67, 163–173.

28. Karoutas, A. and Akhtar, A. (2021) Functional mechanisms and abnormalities of the nuclear lamina. Nat Cell Biol, 23, 116–126.

29. Guelen, L., Pagie, L., Brasset, E., Meuleman, W., Faza, M.B., Talhout, W., Eussen, B.H., de Klein, A., Wessels, L., de Laat, W. et al. (2008) Domain organization of human chromosomes revealed by mapping of nuclear lamina interactions. Nature, 453, 948–951.

30. Meuleman, W., Peric-Hupkes, D., Kind, J., Beaudry, J.B., Pagie, L., Kellis, M., Reinders, M., Wessels, L. and van Steensel, B. (2013) Constitutive nuclear lamina-genome interactions are highly conserved and associated with A/T-rich sequence. Genome Res, 23, 270–280.

31. Bian, Q., Khanna, N., Alvikas, J. and Belmont, A.S. (2013) beta-Globin cis-elements determine differential nuclear targeting through epigenetic modifications. J Cell Biol, 203, 767–783.

32. Kind, J., Pagie, L., Ortabozkoyun, H., Boyle, S., de Vries, S.S., Janssen, H., Amendola, M., Nolen, L.D., Bickmore, W.A. and van Steensel, B. (2013) Single-cell dynamics of genome-nuclear lamina interactions. Cell, 153, 178–192.

33. Harr, J.C., Luperchio, T.R., Wong, X., Cohen, E., Wheelan, S.J. and Reddy, K.L. (2015) Directed targeting of chromatin to the nuclear lamina is mediated by chromatin state and A-type lamins. J Cell Biol, 208, 33–52.

34. Ye, Q., Callebaut, I., Pezhman, A., Courvalin, J.C. and Worman, H.J. (1997) Domain-specific interactions of human HP1-type chromodomain proteins and inner nuclear membrane protein LBR. J Biol Chem, 272, 14983–14989.

35. Polioudaki, H., Kourmouli, N., Drosou, V., Bakou, A., Theodoropoulos, P.A., Singh, P.B., Giannakouros, T. and Georgatos, S.D. (2001) Histones H3/H4 form a tight complex with the inner nuclear membrane protein LBR and heterochromatin protein 1. EMBO Rep, 2, 920–925.

36. Poleshko, A., Mansfield, K.M., Burlingame, C.C., Andrake, M.D., Shah, N.R. and Katz, R.A. (2013) The human protein PRR14 tethers heterochromatin to the nuclear lamina during interphase and mitotic exit. Cell Rep, 5, 292–301.

37. Hirano, Y., Hizume, K., Kimura, H., Takeyasu, K., Haraguchi, T. and Hiraoka, Y. (2012) Lamin B receptor recognizes specific modifications of histone H4 in heterochromatin formation. J Biol Chem, 287, 42654–42663.

38. Nagarajan, P., Ge, Z., Sirbu, B., Doughty, C., Agudelo Garcia, P.A., Schlederer, M., Annunziato, A.T., Cortez, D., Kenner, L. and Parthun, M.R. (2013) Histone acetyl transferase 1 is essential for mammalian development, genome stability, and the processing of newly synthesized histones H3 and H4. PLoS Genet, 9, e1003518.

39. Loyola, A., Bonaldi, T., Roche, D., Imhof, A. and Almouzni, G. (2006) PTMs on H3 variants before chromatin assembly potentiate their final epigenetic state. Mol Cell, 24, 309–316.

40. Jasencakova, Z. and Groth, A. (2010) Restoring chromatin after replication: how new and old histone marks come together. Semin Cell Dev Biol, 21, 231–237.

41. Jasencakova, Z., Scharf, A.N., Ask, K., Corpet, A., Imhof, A., Almouzni, G. and Groth, A. (2010) Replication stress interferes with histone recycling and predeposition marking of new histones. Mol Cell, 37, 736–743.

42. Rivera, C., Saavedra, F., Alvarez, F., Díaz-Celis, C., Ugalde, V., Li, J., Forné, I., Gurard-Levin, Z.A., Almouzni, G., Imhof, A. et al. (2015) Methylation of histone H3 lysine 9 occurs during translation. Nucleic Acids Res, 43, 9097–9106.

43. Campos, E.I., Fillingham, J., Li, G., Zheng, H., Voigt, P., Kuo, W.H., Seepany, H., Gao, Z., Day, L.A., Greenblatt, J.F. et al. (2010) The program for processing newly synthesized histones H3.1 and H4. Nat Struct Mol Biol, 17, 1343–1351.

44. Parthun, M.R., Widom, J. and Gottschling, D.E. (1996) The major cytoplasmic histone acetyltransferase in yeast: links to chromatin replication and histone metabolism. Cell, 87, 85–94.

45. Kleff, S., Andrulis, E.D., Anderson, C.W. and Sternglanz, R. (1995) Identification of a gene encoding a yeast histone H4 acetyltransferase. J Biol Chem, 270, 24674–24677.

46. Verreault, A., Kaufman, P.D., Kobayashi, R. and Stillman, B. (1998) Nucleosomal DNA regulates the core-histone-binding subunit of the human Hat1 acetyltransferase. Curr Biol, 8, 96–108.

47. Chang, L., Loranger, S.S., Mizzen, C., Ernst, S.G., Allis, C.D. and Annunziato, A.T. (1997) Histones in transit: cytosolic histone complexes and diacetylation of H4 during nucleosome assembly in human cells. Biochemistry, 36, 469–480.

48. Schneider, J., Bajwa, P., Johnson, F.C., Bhaumik, S.R. and Shilatifard, A. (2006) Rtt109 is required for proper H3K56 acetylation: a chromatin mark associated with the elongating RNA polymerase II. J Biol Chem, 281, 37270–37274.

49. Driscoll, R., Hudson, A. and Jackson, S.P. (2007) Yeast Rtt109 promotes genome stability by acetylating histone H3 on lysine 56. Science, 315, 649–652.

50. Han, J., Zhou, H., Horazdovsky, B., Zhang, K., Xu, R.M. and Zhang, Z. (2007) Rtt109 acetylates histone H3 lysine 56 and functions in DNA replication. Science, 315, 653–655.

51. Han, J., Zhou, H., Li, Z., Xu, R.M. and Zhang, Z. (2007) Acetylation of lysine 56 of histone H3 catalyzed by RTT109 and regulated by ASF1 is required for replisome integrity. J Biol Chem, 282, 28587–28596.

52. Han, J., Zhou, H., Li, Z., Xu, R.M. and Zhang, Z. (2007) The Rtt109-Vps75 histone acetyltransferase complex acetylates non-nucleosomal histone H3. J Biol Chem, 282, 14158–14164.

53. Tsubota, T., Berndsen, C.E., Erkmann, J.A., Smith, C.L., Yang, L., Freitas, M.A., Denu, J.M. and Kaufman, P.D. (2007) Histone H3-K56 acetylation is catalyzed by histone chaperone-dependent complexes. Mol Cell, 25, 703–712.

54. Bazan, J.F. (2008) An old HAT in human p300/CBP and yeast Rtt109. Cell Cycle, 7, 1884–1886.

55. Berndsen, C.E., Tsubota, T., Lindner, S.E., Lee, S., Holton, J.M., Kaufman, P.D., Keck, J.L. and Denu, J.M. (2008) Molecular functions of the histone acetyltransferase chaperone complex Rtt109-Vps75. Nat Struct Mol Biol, 15, 948–956.

56. Chen, C.C., Carson, J.J., Feser, J., Tamburini, B., Zabaronick, S., Linger, J. and Tyler, J.K. (2008) Acetylated lysine 56 on histone H3 drives chromatin assembly after repair and signals for the completion of repair. Cell, 134, 231–243.

57. Fillingham, J., Recht, J., Silva, A.C., Suter, B., Emili, A., Stagljar, I., Krogan, N.J., Allis, C.D., Keogh, M.C. and Greenblatt, J.F. (2008) Chaperone control of the activity and specificity of the histone H3 acetyltransferase Rtt109. Mol Cell Biol, 28, 4342–4353.

58. Das, C., Lucia, M.S., Hansen, K.C. and Tyler, J.K. (2009) CBP/p300-mediated acetylation of histone H3 on lysine 56. Nature, 459, 113–117.

59. Burgess, R.J., Zhou, H., Han, J. and Zhang, Z. (2010) A role for Gcn5 in replication-coupled nucleosome assembly. Mol Cell, 37, 469–480.

60. Hoek, M. and Stillman, B. (2003) Chromatin assembly factor 1 is essential and couples chromatin assembly to DNA replication in vivo. Proc Natl Acad Sci U S A, 100, 12183–12188.

61. Malay, A.D., Umehara, T., Matsubara-Malay, K., Padmanabhan, B. and Yokoyama, S. (2008) Crystal structures of fission yeast histone chaperone Asf1 complexed with the Hip1 B-domain or the Cac2 C terminus. J Biol Chem, 283, 14022–14031.

62. Mello, J.A., Silljé, H.H., Roche, D.M., Kirschner, D.B., Nigg, E.A. and Almouzni, G. (2002) Human Asf1 and CAF-1 interact and synergize in a repair-coupled nucleosome assembly pathway. EMBO Rep, 3, 329–334.

63. Moggs, J.G., Grandi, P., Quivy, J.P., Jónsson, Z.O., Hübscher, U., Becker, P.B. and Almouzni, G. (2000) A CAF-1-PCNA-mediated chromatin assembly pathway triggered by sensing DNA damage. Mol Cell Biol, 20, 1206–1218.

64. Ridgway, P. and Almouzni, G. (2000) CAF-1 and the inheritance of chromatin states: at the crossroads of DNA replication and repair. J Cell Sci, 113 (Pt 15), 2647–2658.

65. Shibahara, K. and Stillman, B. (1999) Replication-dependent marking of DNA by PCNA facilitates CAF-1-coupled inheritance of chromatin. Cell, 96, 575–585.

66. Tyler, J.K., Collins, K.A., Prasad-Sinha, J., Amiott, E., Bulger, M., Harte, P.J., Kobayashi, R. and Kadonaga, J.T. (2001) Interaction between the Drosophila CAF-1 and ASF1 chromatin assembly factors. Mol Cell Biol, 21, 6574–6584.

67. Liu, W.H., Roemer, S.C., Port, A.M. and Churchill, M.E. (2012) CAF-1-induced oligomerization of histones H3/H4 and mutually exclusive interactions with Asf1 guide H3/H4 transitions among histone chaperones and DNA. Nucleic Acids Res, 40, 11229–11239.

68. Winkler, D.D., Zhou, H., Dar, M.A., Zhang, Z. and Luger, K. (2012) Yeast CAF-1 assembles histone (H3-H4)2 tetramers prior to DNA deposition. Nucleic Acids Res, 40, 10139–10149.

69. Agudelo Garcia, P.A., Hoover, M.E., Zhang, P., Nagarajan, P., Freitas, M.A. and Parthun, M.R. (2017) Identification of multiple roles for histone acetyltransferase 1 in replication-coupled chromatin assembly. Nucleic Acids Res, 45, 9319–9335.

70. Buenrostro, J.D., Wu, B., Chang, H.Y. and Greenleaf, W.J. (2015) ATAC-seq: A Method for Assaying Chromatin Accessibility Genome-Wide. Curr Protoc Mol Biol, 109, 21.29.21–21.29.29.

71. Benson, L.J., Gu, Y., Yakovleva, T., Tong, K., Barrows, C., Strack, C.L., Cook, R.G., Mizzen, C.A. and Annunziato, A.T. (2006) Modifications of H3 and H4 during Chromatin Replication, Nucleosome Assembly, and Histone Exchange. Journal of Biological Chemistry, 281, 9287–9296.

72. Annunziato, A.T., Frado, L.L., Seale, R.L. and Woodcock, C.L. (1988) Treatment with sodium butyrate inhibits the complete condensation of interphase chromatin. Chromosoma, 96, 132–138.

73. Annunziato, A.T. and Hansen, J.C. (2000) Role of histone acetylation in the assembly and modulation of chromatin structures. Gene Expr, 9, 37–61.

74. Stewart-Morgan, K.R., Reverón-Gómez, N. and Groth, A. (2019) Transcription Restart Establishes Chromatin Accessibility after DNA Replication. Mol Cell, 75, 408–414.

75. Amendola, M. and van Steensel, B. (2014) Mechanisms and dynamics of nuclear lamina-genome interactions. Curr Opin Cell Biol, 28, 61–68.

76. Solovei, I., Wang, A.S., Thanisch, K., Schmidt, C.S., Krebs, S., Zwerger, M., Cohen, T.V., Devys, D., Foisner, R., Peichl, L. et al. (2013) LBR and lamin A/C sequentially tether peripheral heterochromatin and inversely regulate differentiation. Cell, 152, 584–598.

77. Vergnes, L., Peterfy, M., Bergo, M.O., Young, S.G. and Reue, K. (2004) Lamin B1 is required for mouse development and nuclear integrity. Proc Natl Acad Sci U S A, 101, 10428–10433.

78. Nagarajan, P., Agudelo Garcia, P.A., Iyer, C.C., Popova, L.V., Arnold, W.D. and Parthun, M.R. (2019) Early-onset aging and mitochondrial defects associated with loss of histone acetyltransferase 1 (Hat1). Aging cell, e12992.

79. Jevtic, P., Edens, L.J., Vukovic, L.D. and Levy, D.L. (2014) Sizing and shaping the nucleus: mechanisms and significance. Curr Opin Cell Biol, 28, 16–27.

80. Jevtic, P., Edens, L.J., Li, X., Nguyen, T., Chen, P. and Levy, D.L. (2015) Concentration-dependent Effects of Nuclear Lamins on Nuclear Size in Xenopus and Mammalian Cells. J Biol Chem, 290, 27557–27571.

81. Karoutas, A., Szymanski, W., Rausch, T., Guhathakurta, S., Rog-Zielinska, E.A., Peyronnet, R., Seyfferth, J., Chen, H.R., de Leeuw, R., Herquel, B. et al. (2019) The NSL complex maintains nuclear architecture stability via lamin A/C acetylation. Nat Cell Biol, 21, 1248–1260.

82. Elsherbiny, A. and Dobreva, G. (2021) Epigenetic memory of cell fate commitment. Curr Opin Cell Biol, 69, 80–87.

83. Chen, S., Luperchio, T.R., Wong, X., Doan, E.B., Byrd, A.T., Roy Choudhury, K., Reddy, K.L. and Krangel, M.S. (2018) A Lamina-Associated Domain Border Governs Nuclear Lamina Interactions, Transcription, and Recombination of the Tcrb Locus. Cell Reports, 25, 1729–1740.e1726.

84. Cabianca, D.S., Munoz-Jimenez, C., Kalck, V., Gaidatzis, D., Padeken, J., Seeber, A., Askjaer, P. and Gasser, S.M. (2019) Active chromatin marks drive spatial sequestration of heterochromatin in C. elegans nuclei. Nature, 569, 734–739.

85. Panigrahi, A. and O’Malley, B.W. (2021) Mechanisms of enhancer action: the known and the unknown. Genome Biol, 22, 108.

86. Agudelo Garcia, P.A., Lovejoy, C.M., Nagarajan, P., Park, D., Popova, L.V., Freitas, M.A. and Parthun, M.R. (2020) Histone Acetyltransferase 1 is Required for DNA Replication Fork Function and Stability. J Biol Chem.

87. Agudelo Garcia, P.A., Nagarajan, P. and Parthun, M.R. (2020) Hat1-Dependent Lysine Acetylation Targets Diverse Cellular Functions. J Proteome Res, 19, 1663–1673.

88. Gruber, J.J., Geller, B., Lipchik, A.M., Chen, J., Salahudeen, A.A., Ram, A.N., Ford, J.M., Kuo, C.J. and Snyder, M.P. (2019) HAT1 Coordinates Histone Production and Acetylation via H4 Promoter Binding. Mol Cell, 75, 711–724 e715.

89. Larson, A.G., Elnatan, D., Keenen, M.M., Trnka, M.J., Johnston, J.B., Burlingame, A.L., Agard, D.A., Redding, S. and Narlikar, G.J. (2017) Liquid droplet formation by HP1alpha suggests a role for phase separation in heterochromatin. Nature, 547, 236–240.

90. Strom, A.R., Emelyanov, A.V., Mir, M., Fyodorov, D.V., Darzacq, X. and Karpen, G.H. (2017) Phase separation drives heterochromatin domain formation. Nature, 547, 241–245.

91. Gibson, B.A., Doolittle, L.K., Schneider, M.W.G., Jensen, L.E., Gamarra, N., Henry, L., Gerlich, D.W., Redding, S. and Rosen, M.K. (2019) Organization of Chromatin by Intrinsic and Regulated Phase Separation. Cell, 179, 470–484.e421.

92. Wang, L., Gao, Y., Zheng, X., Liu, C., Dong, S., Li, R., Zhang, G., Wei, Y., Qu, H., Li, Y. et al. (2019) Histone Modifications Regulate Chromatin Compartmentalization by Contributing to a Phase Separation Mechanism. Molecular Cell, 76, 646–659.e646.

93. Erdel, F., Rademacher, A., Vlijm, R., Tunnermann, J., Frank, L., Weinmann, R., Schweigert, E., Yserentant, K., Hummert, J., Bauer, C. et al. (2020) Mouse Heterochromatin Adopts Digital Compaction States without Showing Hallmarks of HP1-Driven Liquid-Liquid Phase Separation. Mol Cell, 78, 236–249 e237.

94. Gel, B. and Serra, E. (2017) karyoploteR: an R/Bioconductor package to plot customizable genomes displaying arbitrary data. Bioinformatics, 33, 3088–3090.

95. Wickham, H. (2016). Springer-Verlag, New York.

96. Costantini, M., Cammarano, R. and Bernardi, G. (2009) The evolution of isochore patterns in vertebrate genomes. BMC Genomics, 10, 146.

97. Peric-Hupkes, D., Meuleman, W., Pagie, L., Bruggeman, S.W., Solovei, I., Brugman, W., Gräf, S., Flicek, P., Kerkhoven, R.M., van Lohuizen, M. et al. (2010) Molecular maps of the reorganization of genome-nuclear lamina interactions during differentiation. Mol Cell, 38, 603–613.

98. Nagarajan, P., Agudelo Garcia, P.A., Iyer, C.C., Popova, L.V., Arnold, W.D. and Parthun, M.R. (2019) Early-onset aging and mitochondrial defects associated with loss of histone acetyltransferase 1 (Hat1). Aging Cell, 18, e12992.

