## Supplementary Material for "Epigenetic Regulation of Nuclear Lamina-Associated Heterochromatin by HAT1 and the Acetylation of Newly Synthesized Histones"

**Supplementary Information**

**Supplementary Methods**

### Histone Sample Preparation for Epiprofile

Nuclei were isolated and histones extracted as follows. Briefly, 2 x $10^7$ cells were harvested and fractionated into cytoplasmic proteins and nuclei with lysis buffer 1 (LB1 – 50 mM Tris-HCl, pH 7.5, 20 mM NaCl, 1 mM EDTA, 1% Glycerol (vol/vol), 0.5% NP-40 (vol/vol), 0.25% Triton X-100 (vol/vol), supplemented with 1x Roche cOmplete protease tablet, 2 mM Sodium Vanadate, 10 mM Sodium Butyrate, 1 mM DTT) incubated with rotation for 15 min at 4 °C and spun down at 400 *xg* for 5 min at 4 °C. Nuclei were washed with lysis buffer 2 (LB2 – 50 mM Tris-HCl, pH 7.5, 20 mM NaCl, 1 mM EDTA, 0.5 mM EGTA, supplemented with 1x Roche cOmplete protease tablet, 2 mM Sodium Vanadate, 10 mM Sodium Butyrate, 1 mM DTT), incubated and spun as before. Nuclear pellets were suspended in 0.4N $H_2SO_4$ and incubated for 2 h over ice with intermittent vortexing. Cellular debris was pelleted at >20000 *xg* for 15 min at 4 °C, supernatant transferred to 2 mL tube and histones precipitated by addition of ice-cold acetone and incubation O/N at -20 °C.

Samples were washed with ice-cold acetone twice prior to suspension in ABC (100 mM) and quantification via Bradford. Histones were prepared for bottom-up DIA as follows. Briefly, histone proteins (100 µg) were incubated for 15 min at RT with ACN:PA (3: 1 vol/vol) four times to ensure complete conversion of unmodified and mono-methylated lysine residues. Histones were digested with trypsin (1:25 wt/wt) 400 rpm at 37 °C O/N. Histone peptides were incubated with ACN:PA twice to convert all of the newly formed N-termini, dried in a vacuum concentrator and quantified via Nanodrop to estimate concentration and yield prior to mass spectrometry analysis.

**Data-Independent Acquisition Tandem Mass Spectrometry (DIA-MS/MS)**

Histone peptides (1200 ng) were separated with a PepMap C18 column (Thermo Fisher Scientific Cat No ES800A 3 µm, 100 Å, 75 µm x 15 cm) coupled to a Q-Exactive Plus Hybrid Quadrupole Orbitrap mass spectrometer (Thermo Fisher Scientific) by increasing 300 nL/min Buffer B (0.1% FA, ACN) over 75 min as follows: desalt/load peptides on trap 0-5 min 2% B, 5-45 min 45% B, 45-52 min 55% B, 52-56 min 95% B, 56-64 min hold at 95% B, equilibrate to 2% B for 64-75 min. Spectra were collected in data-independent acquisition (DIA) mode with the following parameters: MS1- resolution 35k, AGC target 3e6, maximum fill time 200 ms over a scan range 300-1100 *m/z,* MS2 – resolution 17.5k, AGC target 1e6 with a loop count of 8 and a sliding isolation window of 50 *m/z*. Raw data files were analyzed with EpiProfile 2.0 [194]. Significance was determined with Welch's t-test.

**Data Availability**

RAW mass spectrometry proteomics data reported in this paper has been deposited to PRIDE via massIVE and can be downloaded through the mass spectrometry interactive virtual environment (massIVe) FTP (ftp://massive.ucsd.edu/MSV000087497/) or the accession number (PXD026232) through PRIDE. Further, processed mzML, mztab and the search engine results file generated during the database search for this paper are also deposited with PXD026232.

**Supplementary Figure Legends**

**Supplementary Figure 1. HAT1-dependent chromatin accessibility sites localize to heterochromatic regions of the genome.** A. Pie chart displaying the distribution of HAT1-dependent sites of chromatin accessibility to the indicated genomic features. B. HAT1-dependent sites of chromatin accessibility were localized to the indicated GC-content isochores. C. Locations of HAT1-dependent sites of chromatin accessibility, indicated as vertical lines are shown above a karyoplot of chromosome 1. Below the karyoplot is a graph of gene density on chromosome 1.

**Supplementary Figure 2. Chromatin accessibility patterns are similar between primary and immortalized MEFs.** A. Sliding window averages of log2 fold change of ATAC-Seq data from primary and immortalized $HAT1^{+/+}$ and $HAT1^{-/-}$ MEFs. B. Genome browser view of an 8 Mb region of chromosome 6 showing ATAC-Seq data from primary and immortalized $HAT1^{+/+}$ and $HAT1^{-/-}$ MEFs.

**Supplementary Figure 3. Certain histone modifications are enriched in absence of HAT1.** A. The abundance of peptides containing the indicated modification on histone H3 (top) and histone H4 (bottom) in $HAT1^{+/+}$ and $HAT1^{-/-}$ MEFs (n=4 for each genotype) is plotted as relative abundance (left) or as enrichment/depletion in $HAT1^{-/-}$ cells. B. Histones were isolated from two independent HAT1+/+ and HAT1-/- MEF cell lines and Western blots were probed with the indicated antibodies. The difference in the abundance of H3 K9me2 and K9me3 was quantitated relative to total histone H3.

**Supplementary Table 1. ATAC-Seq sites of differential accessibility in $HAT1^{+/+}$ and $HAT1^{-/-}$ MEFs.**

**Supplementary Table 2. Location of computationally derived HADs in pMEFs.**

**A**

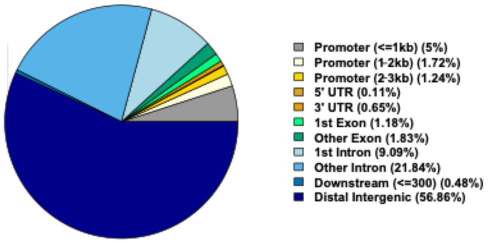


**B**

| ISOCHORE | GC CONTENT | HAT1-DEPENDENT SITES (%) |
|---|---|---|
| L1 | <37% | 16 |
| L2 | 37-41% | 49 |
| H1 | 41-46% | 25 |
| H2 | 46-53% | 8 |
| H3 | >53% | 0 |

**C**

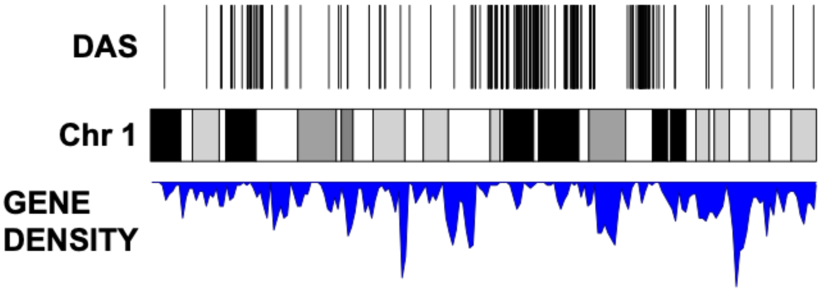


**Supplementary Figure 1**

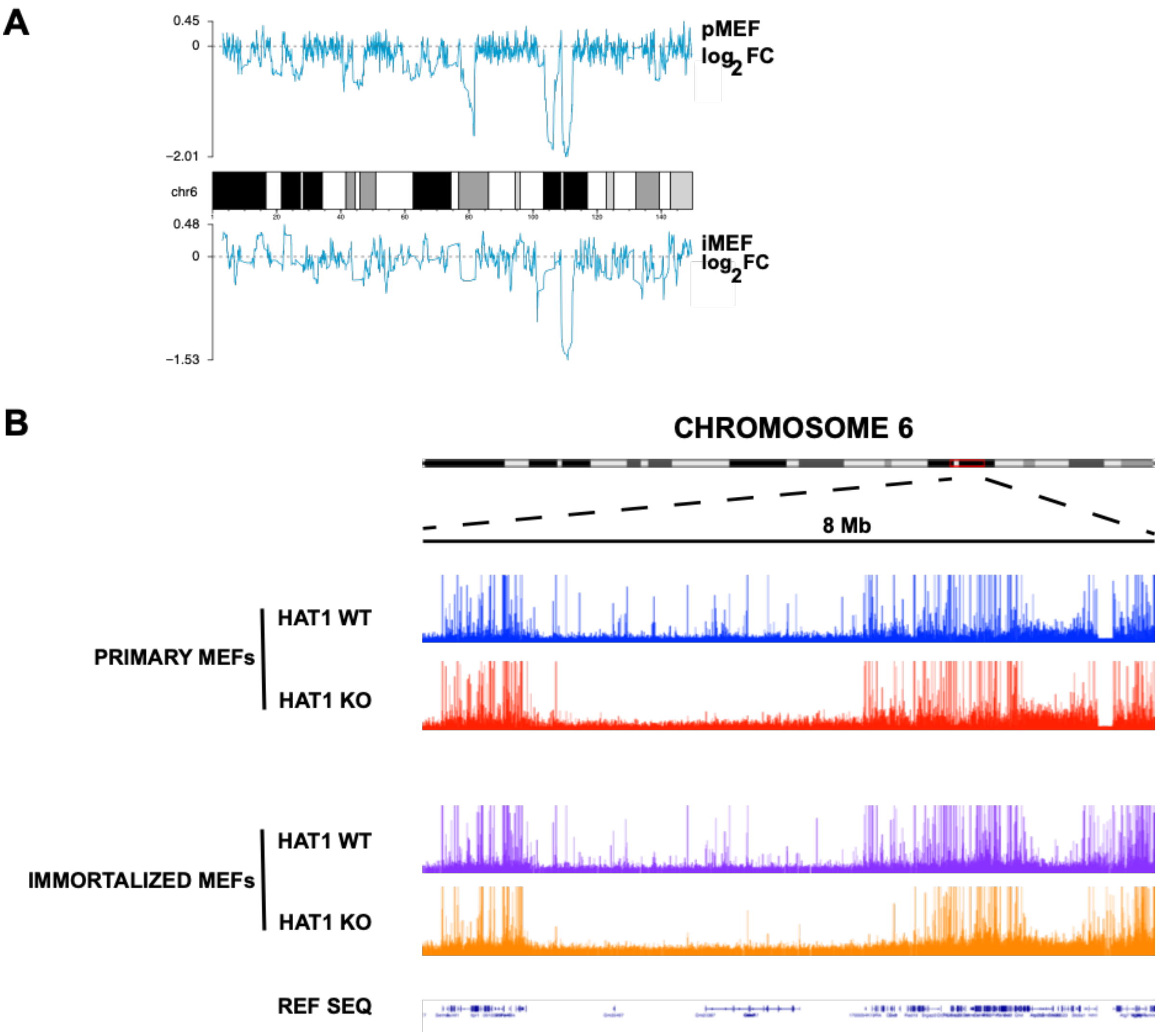


**Supplementary Figure 2**

A

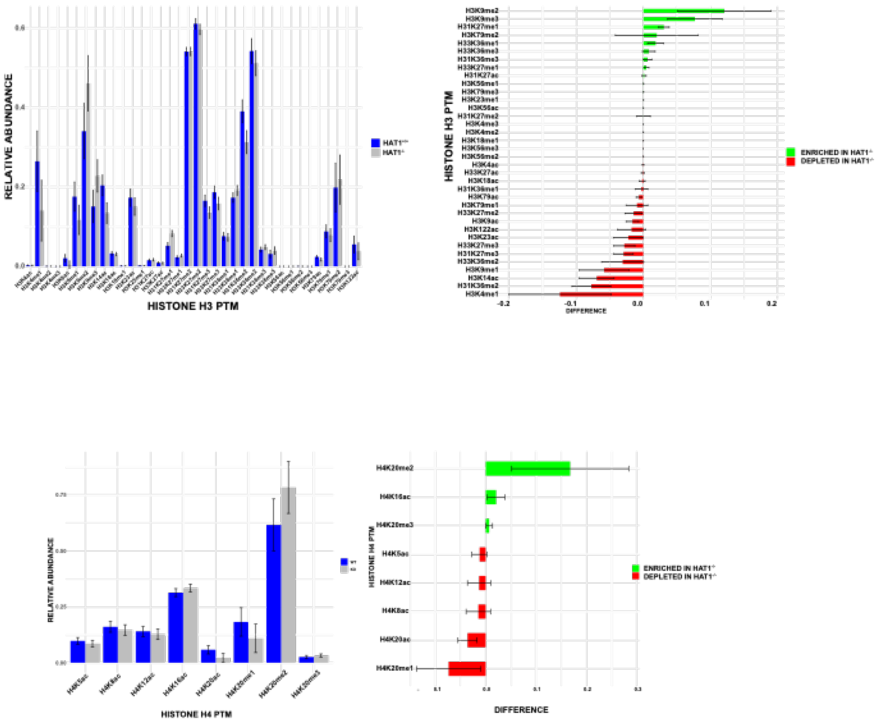


B

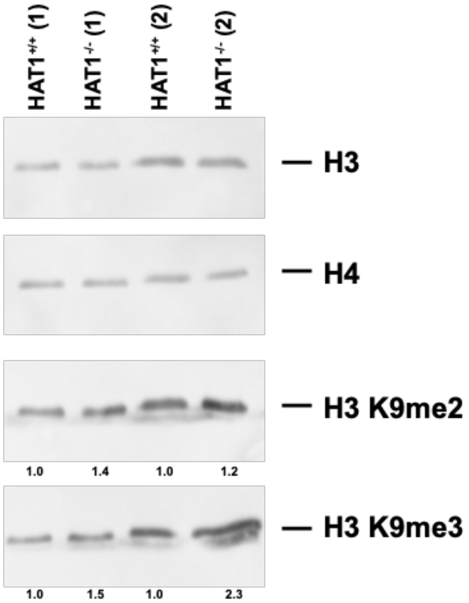


### Supplementary Figure 3
